# Fibrosis resolution in the mouse liver: role of Mmp12 and potential role of Calpain 1/2

**DOI:** 10.1101/2022.02.15.480540

**Authors:** Toshifumi Sato, Kimberly Z. Head, Jiang Li, Christine E. Dolin, Daniel Wilkey, Nolan Skirtich, Dylan D. McCreary, Sylvia Liu, Juliane I Beier, Ryan M. McEnaney, Michael L Merchant, Gavin E Arteel

## Abstract

Although most work has focused on resolution of collagen ECM, fibrosis resolution involves changes to several ECM proteins. The purpose of the current study was two-fold: 1) to examine the role of MMP12 and elastin; and 2) to investigate the changes in degraded proteins in plasma (i.e., the “degradome”) in a preclinical model of fibrosis resolution. Fibrosis was induced by 4 weeks carbon tetrachloride (CCl_4_) exposure, and recovery was monitored for an additional 4 weeks. Some mice were treated with daily MMP12 inhibitor (MMP408) during the resolution phase. Liver injury and fibrosis was monitored by clinical chemistry, histology and gene expression. The release of degraded ECM peptides in the plasma was analyzed using by 1D-LC-MS/MS, coupled with PEAKS Studio (v10) peptide identification. Hepatic fibrosis and liver injury rapidly resolved in this mouse model. However, some collagen fibrils were still present 28d after cessation of CCl_4_. Despite this persistent collagen presence, expression of canonical markers of fibrosis were also normalized. The inhibition of MMP12 dramatically delayed fibrosis resolution under these conditions. LC-MS/MS analysis identified that several proteins were being degraded even at late stages of fibrosis resolution. Calpains 1/2 were identified as potential new proteases involved in fibrosis resolution. CONCLUSION. The results of this study indicate that remodeling of the liver during recovery from fibrosis is a complex and highly coordinated process that extends well beyond the degradation of the collagenous scar. These results also indicate that analysis of the plasma degradome may yield new insight into the mechanisms of fibrosis recovery, and by extension, new “theragnostic” targets. Lastly, a novel potential role for calpain activation in the degradation and turnover of proteins was identified.

---

Regardless of etiology, hepatic fibrosis is an almost universal endpoint of chronic liver injury, when coupled with impaired/incomplete recovery from said injury (1). The clinical sequelae of severe fibrosis/cirrhosis significantly and negatively impact patient survival and quality of life. Given these clinical implications, the mechanisms of hepatic fibrosis initiation and progression have been studied for decades with the goal of identifying better means to prevent and treat this disease (2). Although much progress has been made in our understanding, the translational value of these studies has been hampered by the fact that clinical diagnosis of liver disease is often late-stage, where mechanism-based therapies appear to be less effective.

It has been known that for several years that hepatic fibrosis may resolve, if the underlying causal factor is successfully removed (3). This phenomenon was consistently observed in animal models of fibrosis, but until recently only in rare cases in humans (4). The recent advance of highly effective disease-modifying agents (e.g., direct acting antivirals for hepatitis C viral infection) have dramatically increased the opportunity for fibrosis resolution to occur (5). However, not all patients for achieve fibrosis resolution when the underlying disease has been effectively treated (6). Moreover, in contrast to the development of fibrogenesis, the mechanisms by which fibrosis resolution is mediated are poorly understood and may not share complete mechanistic overlap with fibrosis progression (4). To address both this clinical need and the gap in our understanding, more investigations in this area need to be performed.

The hepatic extracellular matrix (ECM) consists of a diverse range of components that work bi-directionally with surrounding cells to create a dynamic and responsive microenvironment that regulates cell and tissue homeostasis. Under normal conditions, the ECM assists in maintaining organ homeostasis and appropriate responses to injury/stress by dynamically responding to these insults. However, failure to properly regulate these responses can lead to qualitative and/or quantitative ECM changes that are maladaptive (7). Hepatic fibrosis is a canonical example of ECM dyshomeostasis, leading to accumulation of fibrillary ECM, such as collagen, which is considered the pathologic hallmark of fibrosis. Although hepatic fibrosis is considered almost synonymous with collagen accumulation (8), the qualitative and quantitative alterations to the hepatic ECM during fibrosis are much more diverse (9). Indeed, hepatic fibrosis results in qualitative and quantitative changes to almost all components of the hepatic ECM (9). This understanding has let previous work by others to investigate the roles of non-collagen ECM components, as well as enzymes that metabolize them (e.g., matrix metalloproteinases; MMPs) in the initiation and development of experimental and clinical fibrosis (10, 11). In contrast, previous studies of fibrosis resolution have focused almost exclusively on secretion and turnover of collagens and on collagen microstructure (12). The purpose of the current study was to investigate turnover of hepatic ECM during fibrosis resolution in mice.

## Experimental Procedures

Please see Supplemental Materials for additional details on Experimental Procedures.

### Animals and Treatments

8 weeks old, male C57Bl/6J mice purchased from Jackson Laboratory (Bar Harbor, ME) were housed in a pathogen-free barrier facility accredited by the Association for Assessment and Accreditation of Laboratory Animal Care, and procedures were approved by the local Institutional Animal Care and Use Committee. Animals were allowed standard laboratory chow and water *ad libitum*. Mice were administered CCl_4_ (1 ml/kg i.p.; diluted 1:4 in olive oil; Sigma-Aldrich, St. Louis, MO) or vehicle 2×/wk for 4 wks; animals were sacrificed 1, 3, 7, 14 and 28 days after final CCl_4_ injection (Figure 1A). Animals were administered an MMP-12 inhibitor (MMP-408; 10 mg/kg, i.g.; Merck Millipore, Burlington, MA) or vehicle (saline) daily starting after the last injection of CCl_4_. At the time of sacrifice, animals were anesthetized with ketamine/xylazine (100/15 mg/kg, i.p.). Blood was collected from the vena cava just prior to sacrifice by exsanguination and citrated plasma was stored at -80°C for further analysis. Portions of liver tissue were frozen immediately in liquid nitrogen, while others were fixed in 10% neutral buffered formalin or embedded in frozen specimen medium (Tissue-Tek OCT compound, Sakura Finetek, Torrance, CA) for subsequent sectioning and mounting on microscope slides.

**Figure 1.**
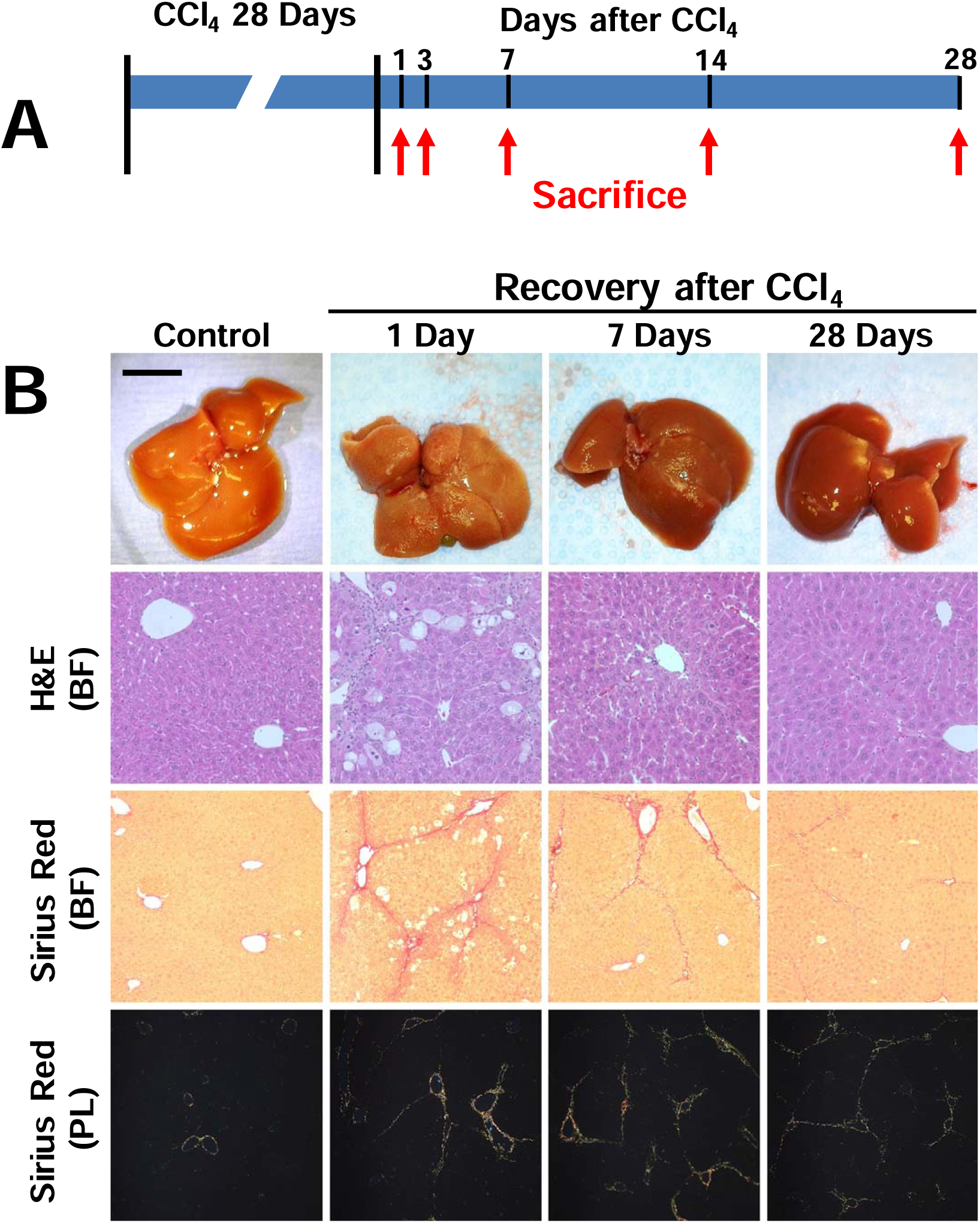
Recovery from liver injury and fibrosis. Panel A: Mice were administered CCl_4_ for 4 weeks to induce hepatic fibrosis as described in Materials and Methods. Animals were sacrificed 1, 3, 7, 14 and 28 days after cessation of CCl4 to track fibrosis resolution. Panel B: Representative photos of livers and photomicrographs (100×) of stained liver sections are shown. Hematoxylin and eosin (H&E) shows general histology, while Sirius red brightfield (BF) and polarized light (PL) show fibrosis.

### Biochemical assays and histology

Plasma levels of Aspartate transaminase (AST) and alanine amino transferase (ALT) were determined spectrophotometrically using standard kits (Thermo Fisher Scientific, Waltham, MA), as described previously (13). Formalin-fixed, paraffin embedded sections were cut at 5 μm and mounted on glass slides. Deparaffinized sections stained with hematoxylin and eosin and pathology was assessed in a blinded manner. ECM accumulation in liver sections was determined by staining with Sirius red/fast green and were visualized via brightfield and polarized light (14). Bright field images of Sirius red staining was quantitated by image analysis, as described previously (13). Specifically, a Molecular Devices (Sunnyvale, CA) Image-1/AT image acquisition and analysis system incorporating an Axioskop 50 microscope (Carl Zeiss Inc., Thornwood, NY) was used to capture and analyze five nonoverlapping fields per section at 100× final magnification. Image analysis was performed using modifications of techniques described previously. Detection thresholds were set for the red color based on an intensely labeled point and a default color threshold. The degree of labeling in each section was determined from the area within the color range divided by the total area.

### Hepatic calpain activity

Calpain activity was measured in liver lysate, using N-succinyl-Leucine-Leucine-Valine-Tyrosine-7-amino-4-methylcoumarin (Suc-Leu-Leu-Val-Tyr-AMC; Supplemental Table 3) was used as the calpain substrate, as described previously (15). Forty μl of liver lysate samples (0.25 mg protein/ml in imidazole buffer containing 63 mM imidazole, 10 mM mercaptoethanol, 1 mM EDTA, 10 mM EGTA, pH 7.3) were incubated at 37 for 1hr with 160 μl of imidazole buffer containing 50 μM Suc-Leu-Leu-Val-Tyr-AMC (dissolved in DMSO). The formation of the cleavage product, 7-amino-4-methylcoumarin (AMC), was measured by fluorometrically (Spectra Max M5, Molecular Devices Corporation) at 360 nm excitation and 430 nm emission wavelengths. Standard curves were generated with free AMC for each experiment and results were normalized to total protein.

### Determination of MMP-2 and 9 by zymography

Zymography was performed as described previously (16). Total hepatic protein was extracted using a lysis buffer consisting of 10 mM cacodylic acid, pH 5.0, containing 150 mM NaCl, 1M ZnCl, 15 mM CaCl_2_, 1.5 mM NaN_3_, and 0.01% Triton X-100. Lysates were then diluted in 2× sample buffer (125 mM Tris HCl, pH 6.8, with 4% SDS, 20% glycerol) without boiling or reduction. The diluted samples were subjected to 0.1% gelatin–10 % SDS polyacrylamide gel electrophoresis (Novex, Carlsbad, CA). The gels were washed with 2.5 % Triton X-100 after electrophoresis and then incubated for 48 hours in the reaction buffer (50 mM Tris, 0.1 M glycine, and 0.1 M NaCl, pH 8.0). The gel was stained with Coomassie brilliant blue R-250 (0.1 % Coomassie Brilliant Blue R250, 50% methanol, and 10% acidic acid) for 1hr, destained with 45% methanol and 10% acetic acid for 30 minutes. The results were visualized and densitometric analysis was performed using an iBright imager and onboard software from Invitrogen and normalized to total protein (Waltham, MA).

### Immunoblots

Liver samples were homogenized in RIPA buffer (20 mM Tris/Cl, pH 7.5, 150 mM NaCl, 1 mM EDTA, 1 mM EGTA, 1% [wt/vol] Triton X-100), containing protease and phosphatase inhibitor cocktails (Sigma). Samples were diluted in 2× sample buffer (see above) and loaded onto 10% SDS-polyacrylamide gels (Invitrogen, Waltham, MA), followed by electrophoresis and Western blotting on to PVDF membranes (Hybond P, GE Healthcare Bio Sciences, Pittsburgh, Pennsylvania). Primary antibodies against proteins of interest were compared to GAPDH as a housekeeping gene (see Supplemental Table 3). The results were visualized and densitometric analysis was performed using an iBright imager and onboard software from Invitrogen (Waltham, MA). Protein levels were normalized either to an internal housekeeping gene (GADPH) or, as in the case for α-fodrin, the ratio of cleaved to total α-fodrin protein signal.

### RNA Isolation and Real-Time RT-PCR

RNA extraction and real-time RT-PCR were performed as described previously (16). RNA was extracted immediately following sacrifice from fresh liver samples using RNA Stat60 (Tel-Test, Ambion, Austin, TX) and chloroform. RNA concentrations were determined spectrophotometrically and 1µg of total RNA was reverse transcribed using the QuantiTect Reverse Transcription Kit (Qiagen, Valencia, CA). Real-time RT-PCR was performed using a StepOne real time PCR system (Thermo Fisher Scientific, Grand Island, NY) using the Taqman Universal PCR Master Mix (Life Technologies, Carlsbad, CA). Primers and probes were ordered as commercially available kits (Thermo Fisher Scientific, Grand Island, NY; see Supplemental Table 1). The comparative C_T_ method was used to determine fold differences between the target genes and an endogenous reference gene (18S). Results were reported as copy number (2^-ΔCT^) or fold naïve control (FC; 2^-ΔΔ*C*T^).

### MMP12 activity

MMP-12 assay kit was purchased from AnaSpec (Fremont, CA). Frozen liver tissue (50-75 mg) was homogenized in the provided assay buffer containing 0.1% (v/v) Triton-X 100, and subsequently centrifuged 10,000×G for 15 min. MMP-12 activity was measured by FRET (490_ex_/520_em_) in the resultant supernatant according to the kit’s instructions and calibrated to commercially-available purified MMP-12 (AnnSpec) and normalized to protein levels in the original homogenate.

### Immunofluorescence

For immunofluorescent detection of elastin, sections of liver were blocked with PBS containing 10 % goat serum for 30 min, followed by incubation overnight at 4 °C with Elastin (Abcam 21607 1:50) antibodies in blocking solution. Sections were washed with PBS and incubated for 3 hrs with a secondary antibody conjugated to Alexa 488 (Invitrogen, Carlsbad, CA). Sections were washed with PBS and mounted with mounting medium containing 1.5 μg/ml DAPI (VECTASHIELD w/DAPI; Vectastain, Torrance, CA). Specificity of staining was validated by comparison to negative controls in the absence of primary antibody.

### Multiphoton microscopy imaging

Elastin and collagen content in the samples were assessed using multiphoton microscopy (MPM), as described previously (17). MPM were performed on frozen sections with a nonlinear optical microscope based on a confocal imaging system (FV1000MPE, Olympus, Tokyo, Japan; purchased with grant 1S10RR025676-01 awarded to Dr Simon C. Watkins) using an optical microscope and an external tunable mode-locked Ti:sapphire LASER (Coherent, Palo Alto, CA) set to 790-800 nm, 5.4% intensity. LASER emission and signal detection were through a 25x, XL Plan N 1.05LN.A. water immersion objective lens (Olympus). Detection was synchronous across three channels. Nuclei stained by propidium iodide (Invitrogen) were detected on the RXD4 channel (663-737 nm emission filter). Fibrillar collagen second-harmonic generation was detected on the RXD1 channel (350-450 nm emission filter). Elastin-associated fibrillar green autofluorescence was detected on the RXD2 channel (500-550 nm emission filter). A 4x line Kalman integration filter was applied to reduce stochastic noise. The laser was routed to an acousto-optic modulator (AOM) for power attenuation, then through a dichroic mirror (490 nm) and an objective to the tissue sample. Signals from second harmonic generation (SHG) were collected by the same objective lens and recorded by a photomultiplier tube (PMT).

### Preparation of acid hydrolysates of liver samples and Quantitation of Desmosine Crosslinks

Liver samples were homogenized in RIPA buffer (see above) and autoclaved (3 h) twice, at 20 lb/in.2 (1.36 atm) to remove collagen and other proteins. The residue was treated with 0.1 M NaOH for 30 min at 100 °C, washed until neutral, and lyophilized. The isolated elastin was flushed with nitrogen and hydrolyzed in 6 N HCl under vacuum at 105°C for 24-36 h. The hydrolysates were evaporated to dryness and the residue was dissolved in 0.5-1.0 ml water. Appropriate aliquots were then added to 0.1 M sodium phosphate buffer, pH 7.4, containing phenol red and neutralized with 2 M NaOH. Desmosine content was determined using a commercially available ELISA kit (Cusabio, Houston, TX) and normalized to starting tissue wet weight.

### Plasma peptide purification and high resolution LC-MS analysis

100 µL PBS containing 0.1X iRT standards (Biognosis Inc, Beverly, MA) was added to 100 µL plasma. 200 µL 20% w/v TCA was mixed into each sample on ice. Samples were then be incubated for 1 hour at 4°C. The precipitate was pelleted by centrifugation at 16,000 × G for 10 min at 4°C; pellets were discarded. The centrifugation was then repeated with the supernatant samples, followed by discard of the second pellets. Samples were then desalted and concentrated using solid phase extraction (Waters Oasis HLB µElution 30 µm plate, part no. 186001828BA) as described (18). Solid phase extraction eluates were dried in a SpeedVac, and dried peptides were resuspended in 25 µL 2% v/v acetonitrile/ 0.1% v/v formic acid. Peptide concentrations were measured by absorbance (NanoDrop2000) at 205 nm and concentrations estimated using a plasma tryptic digest standard curve. Peptide samples (0.2 ug) were separated using an 12cm, 360µm OD x 100µm ID fused silica tip packed with Aeris Peptide 3.6µm XB-C18 100Å material (Phenomenex, Torrance, CA, USA). Peptides were eluted under a 90min 2%-44% acetonitrile gradient and transferred by Nanospray Flex nanoelectrospray source (ThermoFisher) into an Orbitrap Elite - ETD mass spectrometer (ThermoFisher) using an Nth Order Double Play (Xcalibur v2.2, ThermoFisher) with FTMS MS1 scans (240,000 resolution) collected from 300-2000m/z and ITMS MS2 scans collected on up to twenty peaks having a minimum signal threshold of 5,000 counts from the MS1 scan event.

### Peptidomic Data Analysis

RAW files were searched in Peaks Studio X (Bioinformatics Solutions Inc., Waterloo, ON, Canada) using the Denovo, PeaksDB, PeaksPTM, and Label Free Q algorithms and the 2/12/2019 version of the UniprotKB reviewed reference proteome canonical and isoform Homo sapiens sequences (Proteome ID UP000005640). Search parameters included: variable methionine, proline and lysine oxidation (+15.9949 Da), no enzyme specified, 15 ppm precursor error for MS1 Orbitrap FTMS data, 0.5 Da error for MS2 data sets, and common PTMs in the PeaksPTM algorithm. The peptide, feature, and protein-peptide csv files were exported from the Label Free Q result for statistical tests in Microsoft Excel 2016 and R v3.5.0. High confidence peptide assignments (PeaksX criteria –logP scores with p-value <0.05) were exported into comma separated values files (.csv) for upload and analysis by the Proteasix (http://www.proteasix.org/) algorithm using the ‘Peptide-centric’ prediction tool based on curated, known and observed cleavage events enabling assignment of protease identities from the MEROPS database as described previously (19) The positive predictive value (PPV) cut-off for the prediction algorithm output was set to 80%. Protein-protein interaction network analysis of regulated proteomic data sets (q-value <0.05) was performed using Search Tool for the Retrieval of Interacting Genes/Proteins, STRING v11 (20), with the highest confidence score (0.900).

### Data Sharing

Acquired LCMS data (.RAW). Uniprot mouse sequence database annotated with Biognosys iRT standards, metadata for sample key and PeaksX search results exported as an excel file (.xlsx) files have been deposited in MassIVE (massive.ucsd.edu/ProteoSAFe/static/massive.jsp) data repository (MassIVE ID: 000089311) with the Center for Computational Mass Spectrometry at the University of California, San Diego.

### Public data mining

The expression of Capn1 and Capn2 in hepatic from fibrosis was explored, using data from public study GSE135251 [GSE135251-1, GSE135251-2] (21, 22), which includes 46, 48, 54, 54 and 14 patients at fibrosis stage from 0 to 4 [by METAVIR score (23)], respectively. Gene expression profile and clinical information was downloaded from GEO database. The gene count data was normalized by R package DESeq2 normTransform function (24). Expression for *CAPN1* and *CAPN2* as a function of fibrosis stages were visualized by violin plot.

## Results

### Recovery from liver injury and fibrosis

All animals survived exposure to CCl_4_ and no morbidity or mortality was reported during the recovery phase. As expected (25), 4 wks CCl_4_ administration caused severe liver injury and hepatic fibrosis in mouse liver as assessed by histology (Figure 1B) and by transaminase release (Figure 2A). Four weeks of CCl_4_ administration increased hepatic expression of key fibrogenic genes (*Col1a1, Acta2, P4htm, Lox* and *Loxl2*) significantly compared with vehicle controls (Figure 2A). Hepatic fibrosis and liver injury rapidly resolved after cessation of CCl_4_ exposure (Figure 1B), with only thin septa of collagen staining observable after 4 wks recovery (Figure 1B). Likewise, transaminase values and expression of key fibrogenic genes all returned to baseline values after 4 weeks of recovery (Figure 2A).

**Figure 2.**
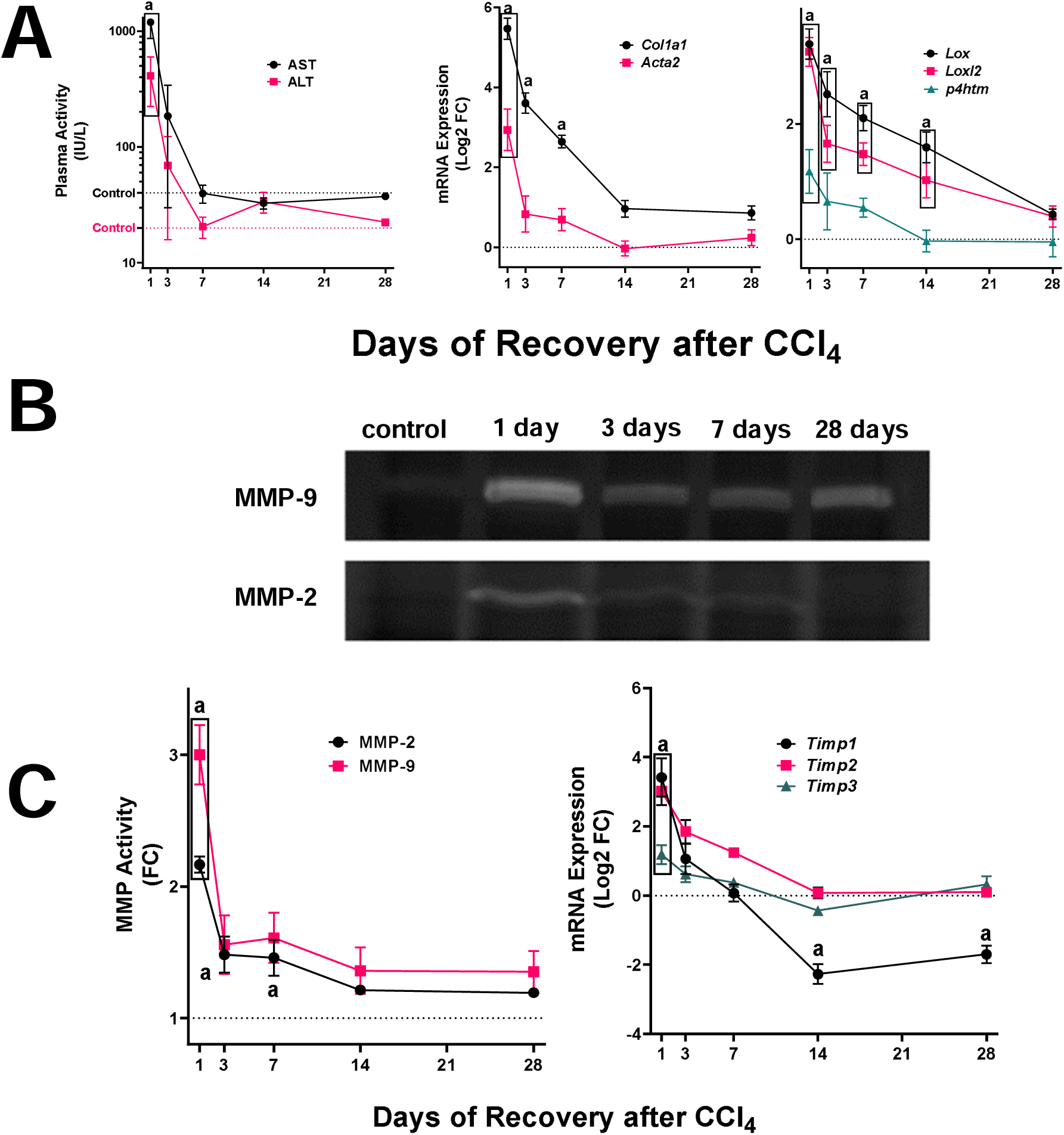
Effect of CCl_4_ and recovery on indices of liver injury and fibrosis. Experimental groups and approaches are as described in Experimental Procedures. Panel A shows plasma transaminase (AST and ALT; IU/L) levels and hepatic mRNA expression of key indices of fibrosis (*Col1a1, Acta2, Lox1, Lox2* and *P4htm*; Log2FC) are shown. Panel B shows representative zymographs demonstrating gelatin lysis by MMP-2 and -9. Panel C shows densitometric analysis of zymography (left; FC) and hepatic expression of key tissue inhibitors of metalloproteinases (*Timp1, Timp2* and *Timp3*; right; Log2FC). Quantitative data are reported as means ± SEM (n=4-6). ^a^*P <*0.05 compared with absence of CCl_4_ by ANOVA using Bonferroni’s post hoc test.

### Effect of CCl_4_ on expression and activity of MMPs

Even in cases where there is a net increase in ECM in the liver (e.g., fibrosis) overall turnover is also increased (26). This turnover is known to be mediated, at least in part, by matrix metalloproteinases (MMPs). 4 wk of CCl_4_ administration increased the expression (Figure 2A) and activity (Figures 2B and 2C) of the collagenases, MMP2 and MMP9. The mRNA expression of inhibitors of these MMPs (i.e., *Timp1, Timp2* and *Timp3*) were also induced with a similar profile (Figure 2C). As was observed with expression of key indices of fibrogenesis (Figure 2A; see above), the increase in expression/activity of these factors largely returned to baseline levels after 28 days of recovery.

In addition to collagen deposition, fibrosis involves quantitative and qualitative changes to a myriad of ECM proteins and MMPs (9, 27). The expression of all hepatic MMPs was therefore determined (Figure 3A). Analogous to findings with *Mmp2* and *Mmp9*, all hepatic MMPs were induced by experimental fibrosis (Figure 3A). The expression pattern of the majority of these MMPs was also similar to *Mmp2* and *Mmp9* and returned to baseline 28 days after the cessation of CCl_4_ (Figure 3A). The one contrast was the expression of *Mmp12*, which was the most robustly induced MMP, both by fold change (1000-fold over control), as well as by copy number (Figure 3A). Unlike other MMPs, expression and protein levels of *Mmp12* were still elevated ∼10-fold over baseline 28 days after cessation of CCl_4_ (Figure 3A).

**Figure 3.**
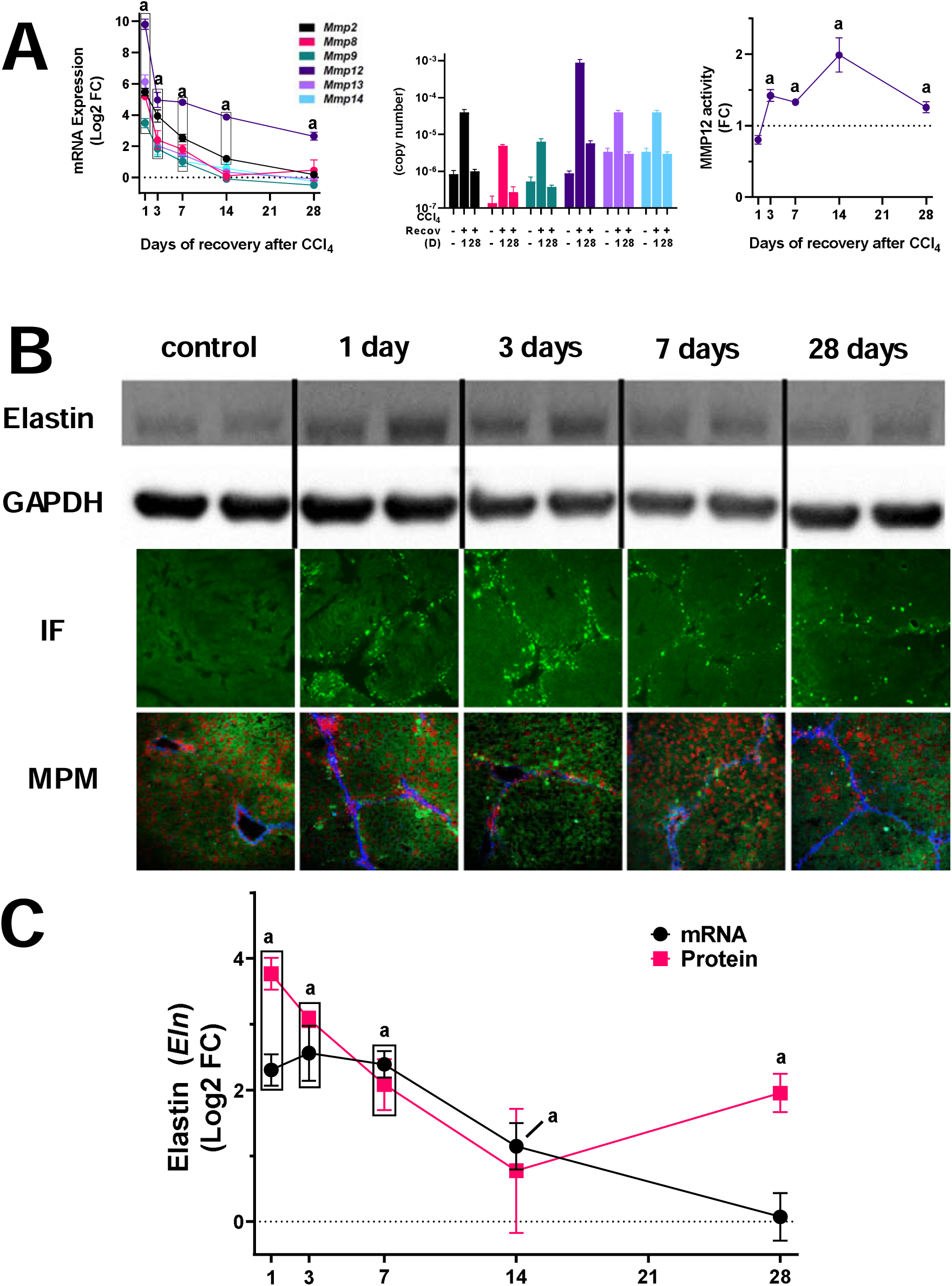
Hepatic expression of MMPs and elastin after fibrosis and recovery. Experimental groups and approaches are as described in Experimental Procedures. Panel A shows hepatic mRNA expression of key indices of fibrosis of all hepatic matrix metalloproteinases (Mmps) both as Log2FC (left) and copy number (right). Panel B shows elastin protein in the liver by Western blot (upper), immunofluorescence (100×; middle) and multiphoton microscopy (MPM, 100×, lower). Elastin immunofluorescence is depicted as intese green staining against the background of hepatic tissue autofluorescence (light green). MPM images depict elastin (green), collagen (blue) and stained nuclei (red). Panel C summarized densitometric quantitation of elastin by Western blot (FC). Quantitative data are reported as means ± SEM (n=4-6). ^a^*P <*0.05 compared with absence of CCl_4_ by ANOVA using Bonferroni’s post hoc test.

### Effect of CCl_4_ on elastin accumulation

Elastin is a key substrate for MMP12 that increases during CCl_4_-induced fibrosis (10). Expression and protein levels of elastin were therefore determined (Figures 3B and C). Although the mRNA expression of elastin (*Eln*) was only moderately (<4 f-fold) induced by 4 weeks of CCl_4_ administration and returned to baseline after 28 days of recovery, protein levels increased ∼15-fold compared with naïve control mice (Figures 3B and 3C). Moreover, elastin ECM was still significantly (∼5 fold) elevated over naïve control values even after 28 days of recovery. As has been observed in previous studies (10), elastin accumulation appeared to correlate with areas of fibrotic changes (Figure 3B). Elastin accumulation appears to center on areas of fibrogenesis and collagen accumulation (see Figure 3B). To investigate this further, the colocalization of elastin and collagen were determined using multi-photon microscopy (Figure 3B), as it allows observation of the elastin and collagen fiber networks simultaneously (17). Elastin (green fluorescence) accumulation was highest on the first day of recovery and was closely associated with collagen (blue fluorescence) signals, consistent with the assessment of elastin immunofluorescence (Figures 3B).

### MMP12 inhibition delays fibrosis resolution

Previous work with knockout mice indicated that MMP12-mediated elastin degradation plays a protective role in the development of CCl_4_-induced fibrogenesis (10); however, the role of MMP12/elastin on resolution of fibrosis was unknown. The effect of the MMP12 inhibition on resolution of fibrosis was therefore determined (Figure 4). CCl_4_-exposed mice were administered the MMP12 inhibitor, MMP-408, daily after cessation of toxicant exposure (see Materials and Methods). MMP-408 did not significantly affect recovery of transaminases after CCl_4_ exposure (Figure 4A). MMP-408 also did not significantly impact the expression of fibrogenic genes (e.g., *Col1A1*) during the recovery phase (Figure 4A). On the other hand, MMP408 impacted the resolution of collagenous fibrosis, as determined by Sirius red staining (Figures 4B and C). Image-analysis and quantification of Sirius red staining indicated that the amount of collagen present in the liver was double 1 wk after cessation of CCl_4_ administration compared to the mice administered the vehicle for MMP408 (Figure 4C). Although this effect of MMP408 was not present after 28 days of recovery, the fibrous septa in the MMP-408 group were far less defined than in the vehicle treated (Figure 4B and inset). The elastin signal in the multiphoton microscopy analysis shows both immature and mature elastin [Figure 3B (17)]. Therefore, it reflects the total amount of elastin accumulation in the liver. In contrast, the crosslinking of mature insoluble elastin is mediated via post-translational incorporation of the amino acid desmosine. Desmosine is unique to mature elastin in mammalian species and can therefore be employed as a surrogate marker of mature elastin (28). The amount of desmosine present in the liver tissues after CCl_4_ was therefore determined by ELISA (Figure 4C), as described in Materials and Methods. Although the total amount of elastin detected in the liver increased by 4 wks of CCl_4_ administration (Figures 3), the desmosine content in the liver actually decreased to ∼50% of levels in naïve mice (Figure 4C). These data suggest that the majority of the new elastin detected during fibrosis is immature uncrosslinked elastin. During the resolution phase desmosine content in livers from mice administered vehicle increased, with values not significantly different that naïve control mice 28 days after cessation of CCl_4_. In contrast, administration of MMP408 completely attenuated the increase in this index of mature elastin (Figure 4C).

**Figure 4.**
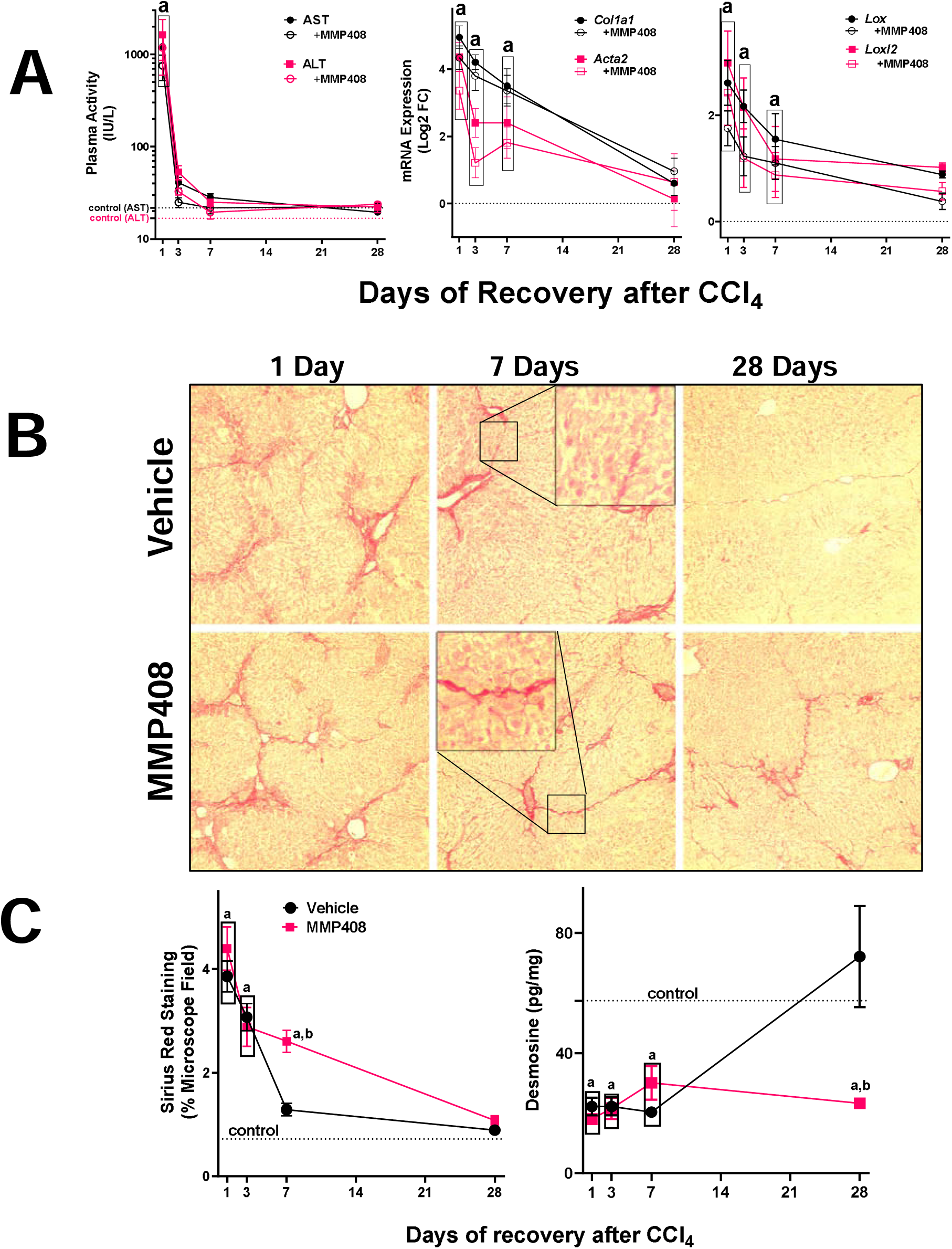
The effect of MMP-408 on recovery from liver injury and fibrosis. Experimental groups and approaches are as described in Experimental Procedures. The effect of MMP-408 on liver transaminase levels (Figure 3A, left) and fibrosis genes (Figure 3A, middle and right) during recovery were determined. The effect of MMP-408 on resolution of fibrosis was assessed by Sirius Red staining (100×) (Figure 3B). Intensity of Sirius red staining was assessed by image analysis (Figure 3C, left). Quantitative data are reported as means ±SEM, (n = 8-10). ^a^*P <*0.05 compared with absence of CCl_4_; ^*b*^*P <*0.05 compared with vehicle group by ANOVA using Bonferroni’s post hoc test.

### The degradome changes dramatically during fibrosis resolution

Resolution of fibrosis requires coordinated remodeling of the liver. The net effect of fibrosis and resolution on hepatic protein degradation has not been previously determined. The influence of CCl_4_ and 4 wks recovery on the plasma peptide “degradome” was therefore determined (Figure 5). Across the groups, 694 peptides were identified, corresponding to 181 distinct parent proteins. The parent proteins that contributed to the pattern of significantly changed peptides were analyzed by StringDB, followed by K-means cluster analysis [Figure 6 (20)]. This database queries physical and functional protein-protein associations that integrates both experimental and predicted associations. The output creates both graphical and categorical enrichment patterns in the queried dataset. Cluster analysis yielded 7 enriched clusters in the total degradome, with a strong enrichment of proteins associated with Gene Ontology (GO) terms for “high-density lipoprotein particle” (GO:0034464; Cluster 1), “collagen trimer” (GO:0005581; cluster 6) and “nucleosome” (GO:0000786; Cluster 7) (Figure 6; Table 1).

**Figure 5.**
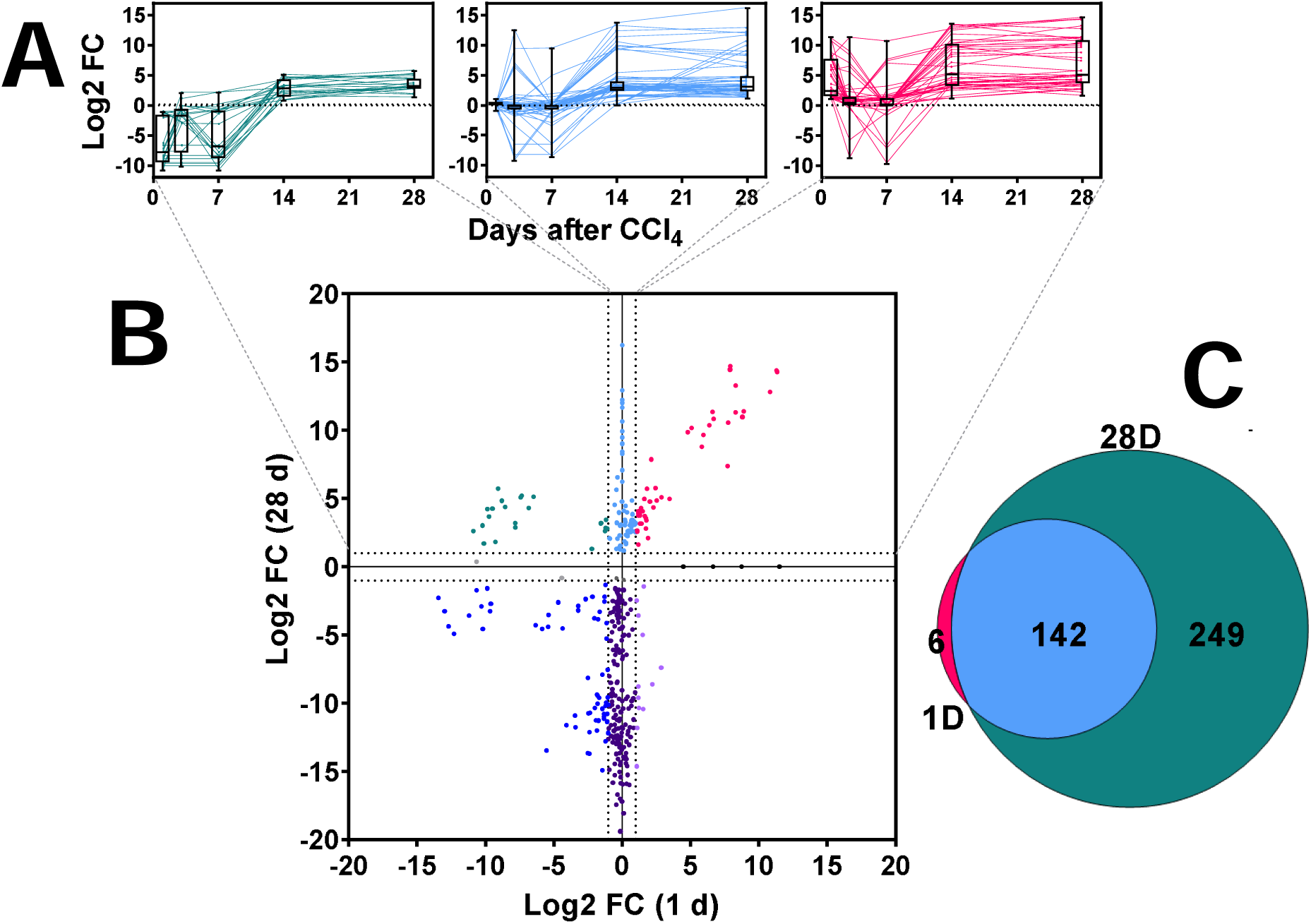
Effect of fibrosis and recovery on the plasma degradome. Experimental groups and approaches are as described in Experimental Procedures. Plasma peptides were measured by LC-MS/MS, as described in experimental procedures and compared naive control mice (Log2FC). Panel A shows the pattern of changes in peptides over the course of the recovery period that were increased at 28 days. Panel B compares the change in peptides after 28 days of recovery, versus those a 1 day. Panel C is a Venn diagram comparing the peptides increased at 28 days of recovery to those increased at 1 day of recovery. Box and whisker plots depict the range, median and interquartile range of the changes of the degraded proteins over the period of recovery (Panel A).

**Figure 6.**
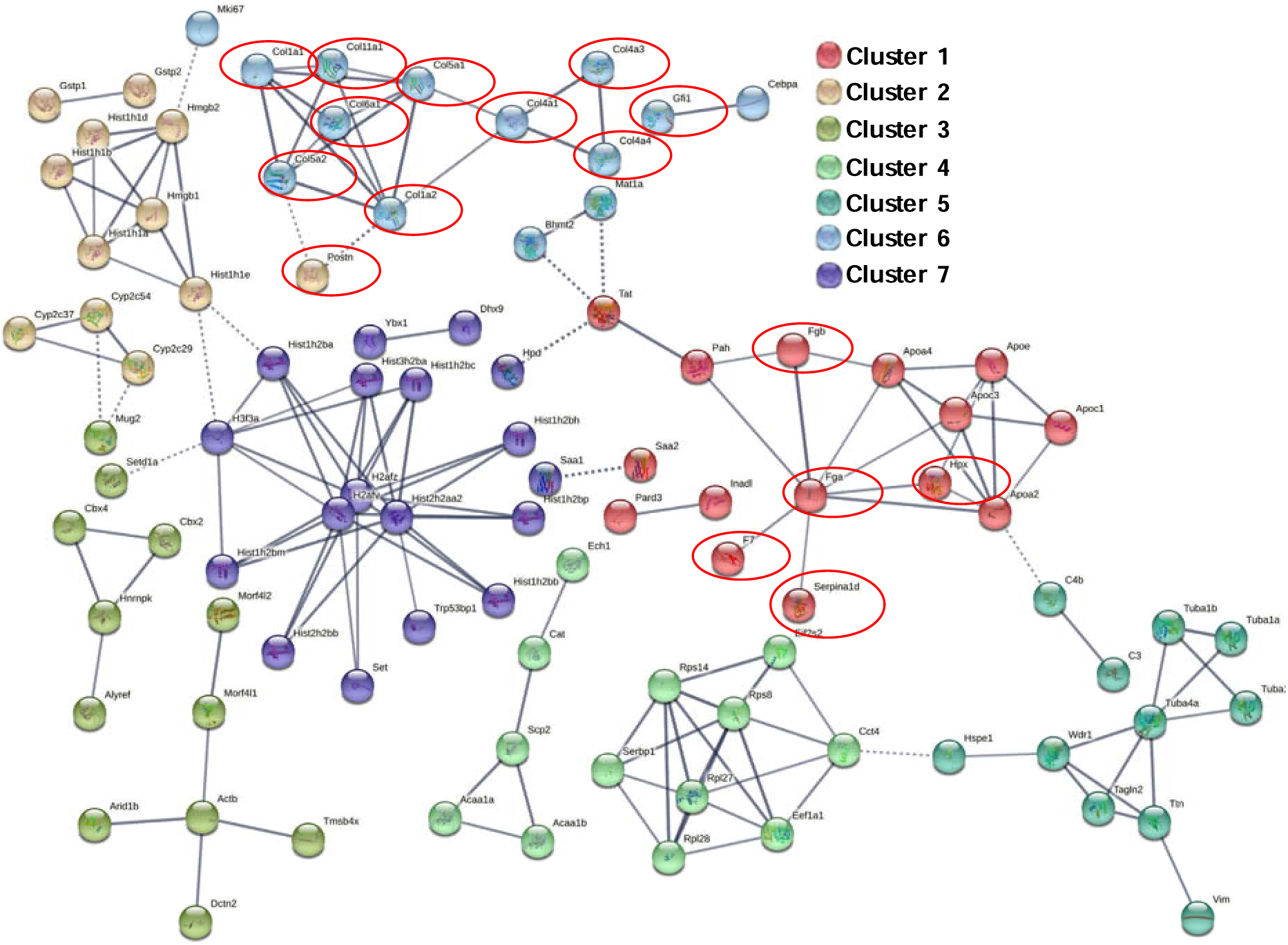
StringDB clustering analysis of peptides significantly increased 28 days after cessation of CCl_4_. Red circled parent proteins are defined as “Matrisome” by the matrisome project (www.matrisomeproject.mit.edu) (55). Cluster groups are summarized in Table 1.

**Table 1.**
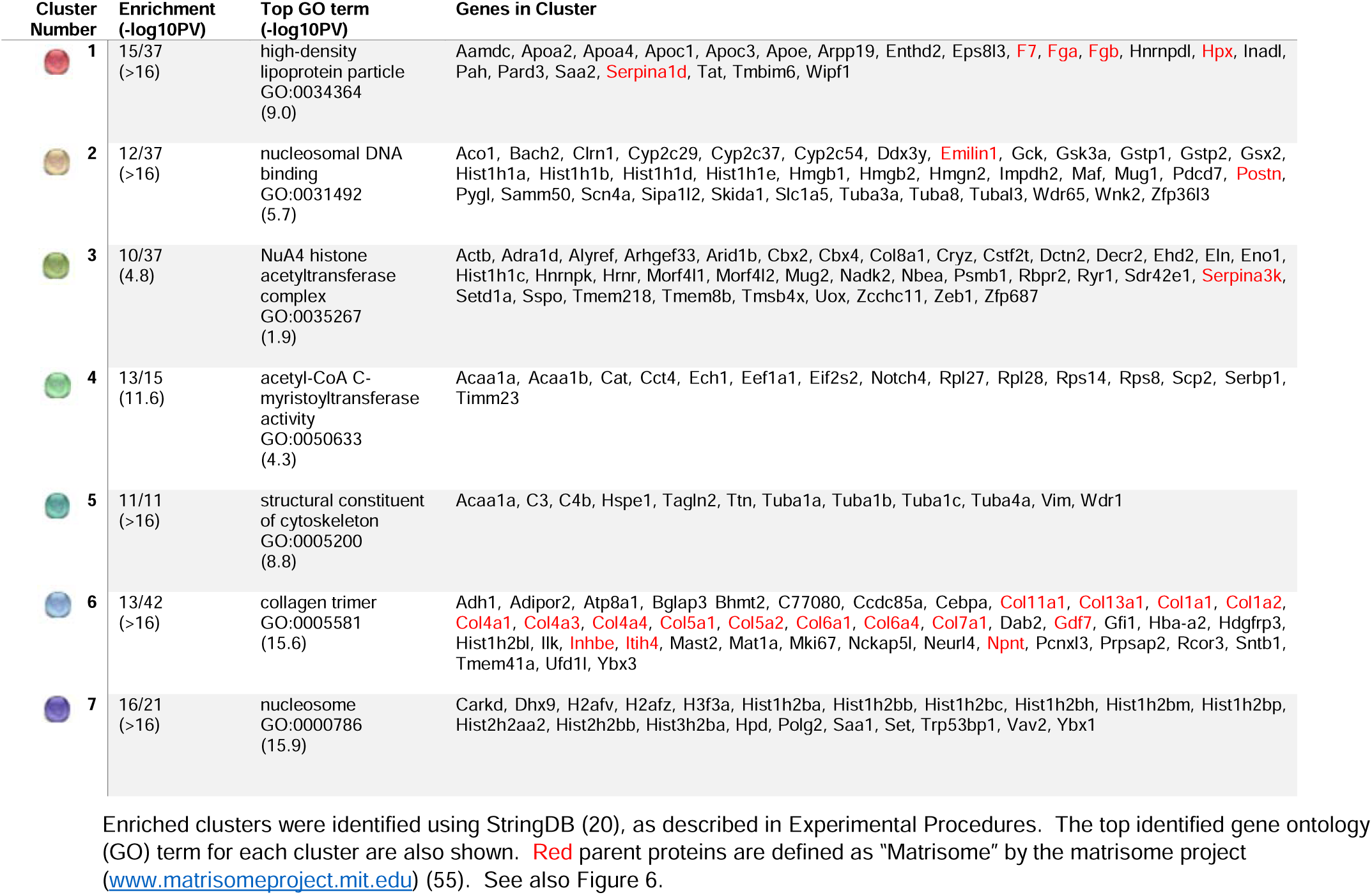
Clusters identified by StringDB to be enriched in the degradome.

The degradome from 1 and 28 D of recovery were compared in more detail to represent the peak of injury (1 D) to near resolution of injury (28 D; Figures 1 and 2). A total of 255 peptides were significantly changed at least >2 fold (compared to naïve control) at these timepoints (Figure 5A, B and C). Although there was considerable overlap between these 2 timepoints (Figure 5C), the amount of peptides altered at the 28 D timepoint was ∼2.5-fold more compared to the 1 D timepoint. A total of 144 peptides, corresponding to 82 distinct proteins, were significantly increased at 1 D after CCl_4_ and/or 28 D after CCl_4_ (Figure 5A and B, Supplemental Table 4). All peptides that significantly increased at 28 D after recovery tended to be at relatively low levels 3 and 7 days after cessation of CCl_4_, then increasing at the 14 D timepoint (Figure 5A).

Many proteases cleave substrates with high specificity with limited cleavage promiscuity. Thus, the fragment sequence of degraded proteins can be informative on proteases that may have generated this pattern. Proteasix is an open-source peptide-centric tool to predict in silico the proteases involved in generating the observed peptide features (29). Figure 7A is a frequency distribution figure comparing the predicted proteases that generated the significantly increased peptides at the 1 and 28 D timepoint (vs. naïve control). The top predicted proteins, calpains 1 and 2 were shared between the timepoints under these conditions (Figure 7A). The expression of these splice variants (*Capn1* and *Capn2*) was determined via real-time rtPCR (Figure 7B). The copy number of *Capn2* was approximately an order of magnitude higher than *Capn1* in liver (3.1×10^−5^ vs. 4.3×10^−6^, respectively) under basal conditions. The expression of both isoform variants was increased by CCl_4_ administration but returned to baseline by 28 D of recovery (Figure 7B). In contrast, predicted enzyme activity of CAPN, both by cleavage of α-fodrin and by enzymatic analysis (Figure 7B), indicated that CAPN activity was elevated at the 28 D timepoint. Several of the predicted CAPN1/2 substrates for these enzymes (Figure 7A) were analogous to the key clusters identified by StringDB analysis for the degradome as a whole (Figure 6 and Table I), such as fibrinogens (cluster 1), collagens (cluster 6) and histones (cluster 7) and their corresponding GO terms.

**Figure 7.**
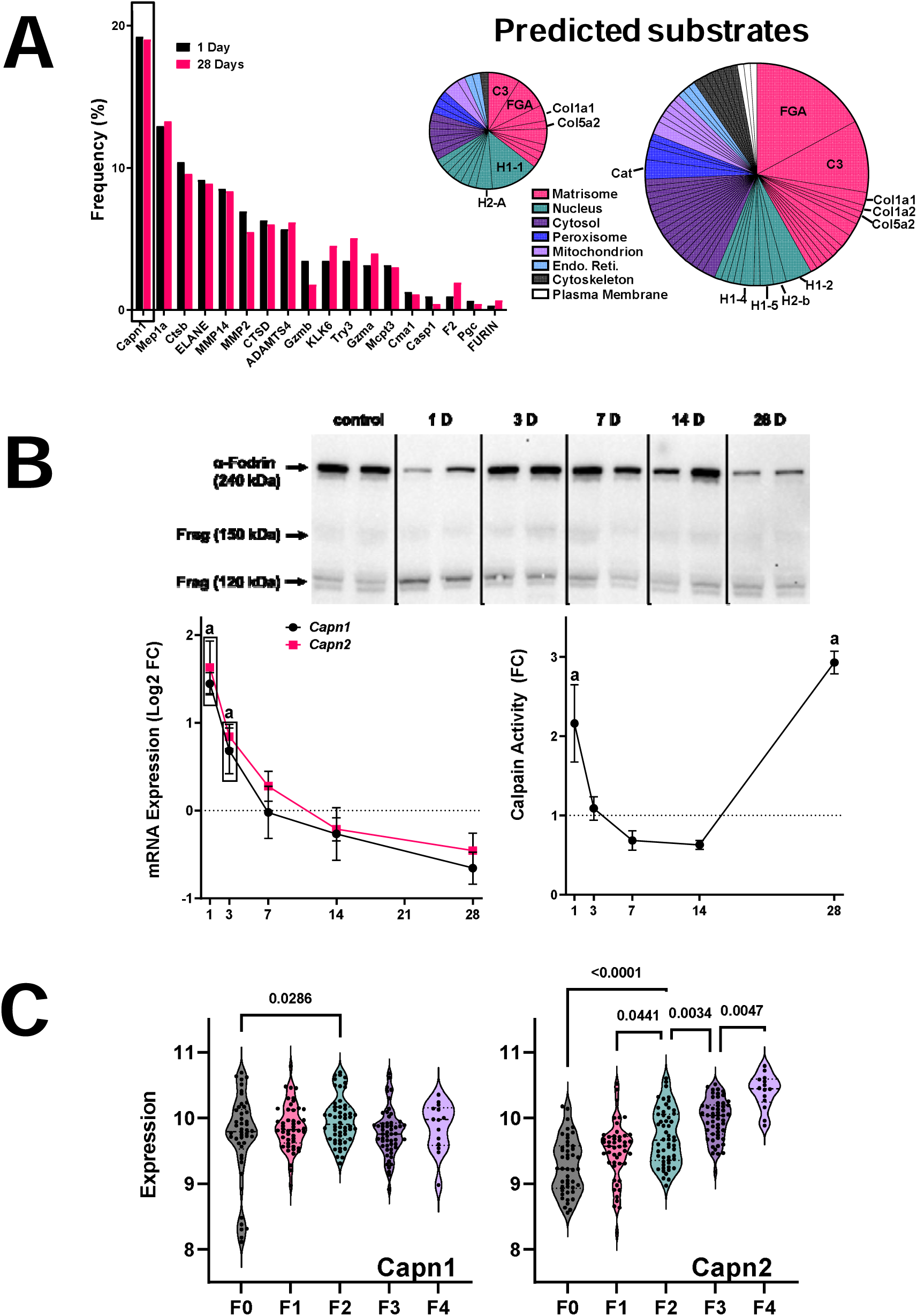
Expression and activity of predicted protease from ProteaSix. Experimental groups and approaches are as described in Experimental Procedures. ProteaSix was used to predict proteases that formed these peptides (Figure 6A). Panel B (upper) depicts western blot analysis of α-fodrin. The 240 kDa band represents intact α-fodrin, while the 150 and 120 kDa bands represent calpain-cleaved fragments. Densitometric analysis is also shown (lower, right). The expression (lower, left) of *Capn1* and *Capn1* were determined by real-time rtPCR. Panel 6C summarizes the type of parent proteins identified a potential substrates for Capn1/2 cleavage by ProteaSix, organized by subcellular localization. The relative size of the pie charts (1 D vs 28 D) is reflective of the amount of peptides identified at each timepoint. Quantitative data are reported as means ±SE, n=4-6. ^a^*P <*0.05 compared with absence of CCl_4_ by ANOVA using Bonferroni’s post hoc test.

### Capn2 is induced in human NASH fibrosis

These data indirectly suggest that calpains may be novel mediators of hepatic fibrosis and resolution. However, the relevance of these findings to human fibrotic liver disease is unclear. A recent study showed that the expression of calpain proteases was induced in human livers with fibrosis caused by HBV (30). Here, the expression of CAPN1 and CAPN2 was determined in hepatic NASH fibrosis (see Experimental Procedures; Figure 7C). The expression of CAPN1 was slightly increased during early (F2) fibrosis; in contrast, the expression of CAPN2 was increased in a step-wise fashion at every fibrosis stage (Figure 7C).

## Discussion

Direct-Acting Antiviral therapy has for all intents and purposes cured the hepatitis C virus (HCV) epidemic in immunocompetent adults. However, one of the most important issues is whether patients with significant HCV-induced fibrosis/cirrhosis will recover from this injury once a sustained virologic response is achieved (31). For example, improvement of liver stiffness in HCV cirrhosis after successful antiviral therapy is reportedly 30-65%, depending on the study referenced (32-36). Although promising at first glance, these data indicate that around half of the HCV-fibrosis/cirrhosis patients with successful viral clearance do not improve in their liver disease. Furthermore, another alarming concern is that the commonly used non-invasive transient elastographic analysis of liver stiffness tends to overpredict fibrosis/cirrhosis regression in HCV, compared to biopsy-based histologic assessment (31, 37). Indeed, despite achievement of viral clearance, histological assessment of patient livers often continues to show lingering inflammation, hepatocyte and ductular reaction and fibrosis (31, 37). It is therefore an unmet clinical need to accurately predict determining factors of fibrosis resolution. Although accumulation of collagen ECM is the *de facto* definition of fibrosis, the changes to the hepatic ECM that happen during both processes are much diverse than those that happen to collagen (9). Moreover, recent studies have suggested that livers that recover from fibrosis do not completely revert to the naive state and are more sensitive to reinduction of fibrosis (12). In toto, although it is understood that hepatic fibrosis resolution involves more than simple clearance of collagen ECM, the process and its mediators are incompletely understood, at best (38).

Previous studies have indicated that MMP12, most likely via mediating elastin ECM accumulation, is key for the development of hepatic fibrosis (10). As in other solid organs, MMP-12 is almost exclusively expressed at high levels in a subset of macrophages in the liver. In previous work Ramachandran et al. (39) identified a subset of Ly6C^low^ macrophages that were enriched in the resolving livers; these macrophages expressed high levels of MMP-12. Previous studies have shown that genetic or pharmacologic inhibition of MMP12 exacerbates the development of experimental fibrosis. Furthermore, prevention of elastin crosslinking protected against experimental fibrosis (40). More recently, Chen et al. (41) demonstrated that elastin fibers degrade slowly during fibrosis resolution, a finding recapitulated in the current study (Figure 3). The authors of that study suggested that cross-linked elastin stabilizes the fibrotic ECM and may thereby protect it from degradation (41). However, whether MMP12 activity mediates resolution of fibrotic ECM was not clear. Here, it was demonstrated that the MMP12 inhibitor, MMP408, delayed resolution of Sirius red-positive areas of the liver (Figure 4), which supports the hypothesis that elastin degradation is key to overall fibrosis resolution. Although elastin protein levels increased (Figure 3), the amount of mature crosslinked elastin (as determined by desmosine content; Figure 4C) actually was decreased by CCl_4_ exposure. Desmosine content did not return to basal levels until after 28 D of recovery. These results suggest that the increase in elastin content in the liver during fibrosis represents less mature (crosslinked) protein oligomers (Figure 8). This finding is similar in models of photoaging in which damaged skin accumulates disorganized elastin fibrils (42). Interestingly, inhibition of MMP12 with MMP408 prevented the recovery of desmosine content during fibrosis resolution, suggesting that MMP12 is critical not only for the degradation of accumulated elastin, but also the reorganization to mature elastin (Figure 8). These findings should be explored in future studies.

**Figure 8.**
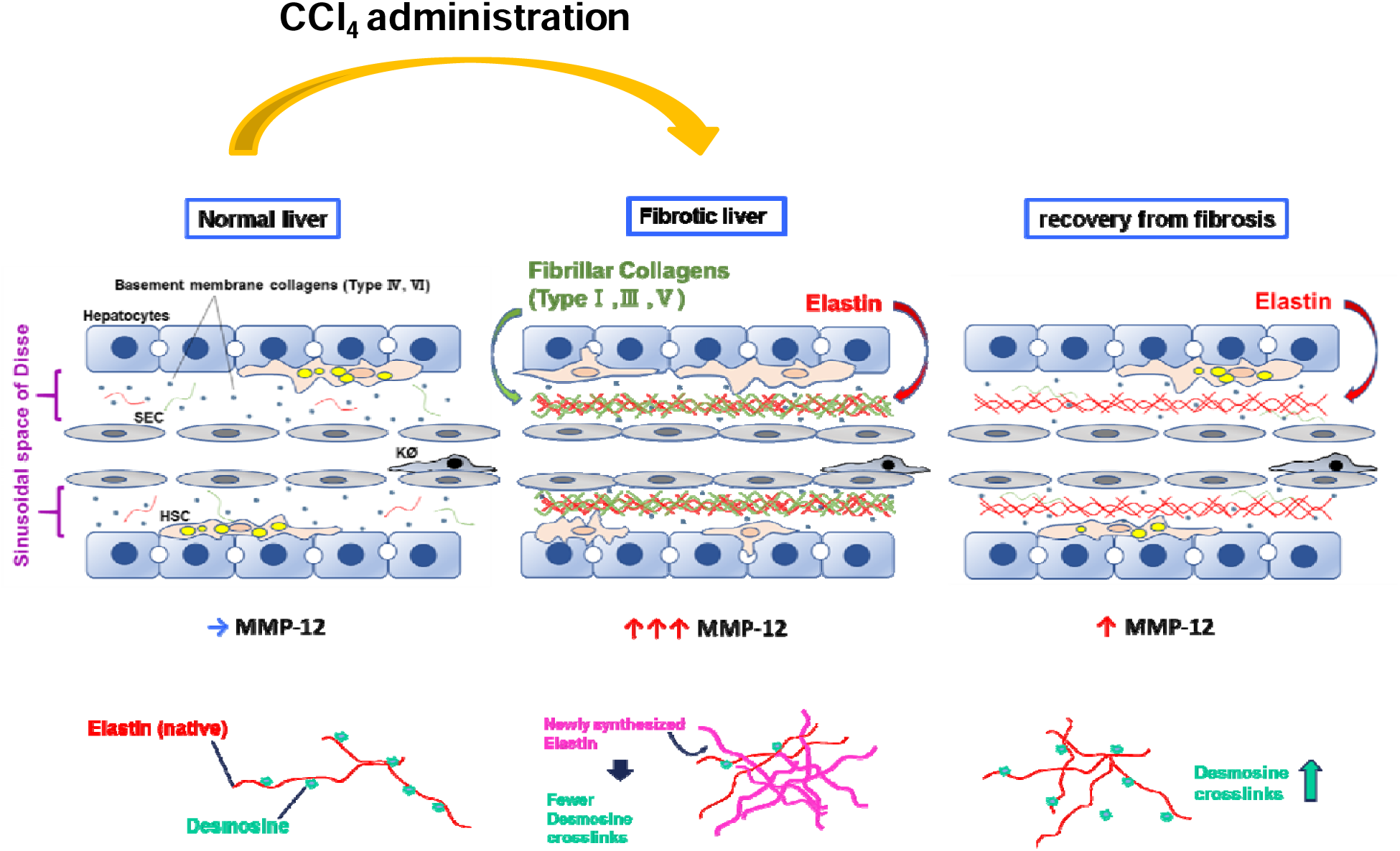
Proposed role of MMP-12 and elastin during recovery from liver fibrosis. Hepatic activity of MMPs was robustly induced; but MMP-12 returned to baseline after recovery from fibrosis. Elastic fibers, which are substrates of MMP-12, are also critical in fibrotic ECM formation, and appeared after CCl_4_ administration. Fibrosis resolved and histology was nearly normal after 28 D recovery, but elastin ECM still remained elevated. Mature tropoelastin chains are connected with desmosine cross-links. Newly synthesized immature elastin lacks this post-translational modification. As recovery progresses, total elastin decreases, but desmosine content increases; interestingly, blocking MMP-12 activity prevented both of these phenomena.

Although fibrosis resolution was delayed by MMP12 inhibition, there were no detectable differences in total sirius red staining between vehicle and MMP12 inhibitor 28 days after the cessation of CCl_4_ intoxication (Figure 4C). However, the pattern of staining in the MMP12 inhibitor group was more diffuse and less well-defined compared to vehicle controls at this timepoint (Figure 4B). These data do not necessarily indicate that MMP12 is equivocal in fibrosis resolution. First, as shown by studying the degradome (see Figures 5-6), there is still significant difference in protein turnover 28 days after cessation of CCl_4_, a time at which the liver is almost histologically normal (Figure 1). Moreover, spontaneous resolution of fibrosis in rodents is much more robust than in humans and by extension, humans are likely more sensitive to delays in fibrosis resolution (12). Although not yet studied in human liver disease, previous work has shown that remodeling in other organ systems (e.g., atherosclerosis) are sensitive to variants in MMP12 activity; for example, a gain-of-function promoter polymorphism in MMP12 is associated with protection against coronary artery disease in diabetes (43).

Hypothesis-driven approaches are key to elucidating mechanism. However, the nature of the approach by definition requires prior knowledge. Discovery-based ‘omic techniques coupled with informatic analyses can yield new information and insight into disease progression and regression. The peptidome is defined as the population of low molecular weight biologically derived peptides (0.5-3 kDa), within the cells and biologic fluids. These compartments contain peptides critical for normal organismal function. The study of the peptidome (i.e., peptidomics’) is a discipline related to proteomics, but with significant methodological and analytical differences (44). For example, as the original structure of the peptides is of interest, samples are not trypsin digested prior to LC-MS/MS analysis, as is the case for bottom-up proteomic approaches. This difference originally limited peptide identification, as the available databases were based on trypsin-digested peptide fragments (45). However, the development of peptidome-specific identification and analysis tools has addressed this concern (46). In addition to synthesized peptides, the peptidome also contains fragments of proteins degraded by normal and/or abnormal processes (i.e., ‘degradome’). These degradomic profiles are conditionally unique to the metabolic/pathologic status of the organism and reflect not only quantitative changes (i.e., more/less parent protein to degrade), but also qualitative changes (i.e., change in the pattern of protease digestion of parent proteins). The latter subset of the peptidome has generated key interest in some areas of human health as possible (surrogate) biomarkers for disease. Here, LC-MS/MS analysis of the degradome profile in plasma during recovery from experimental fibrosis was analyzed to investigate the general pattern of protein turnover/remodeling during recovery, as well as to identify potential new key players.

When the degradome was analyzed, an interesting pattern resolved. Specifically, although livers were almost histologically-normal after 4 weeks of recovery from CCl_4_-induced experimental fibrosis (Figure 1), there was a relative increase in the turnover of proteins observed in the plasma compartment (Figures 5-6), as determined by analysis of the degradome. This finding is in-line with previous observations by this group that hepatic remodeling in response to injury is much more diverse than can be observed histologically (13). In addition to the appearance of degraded ECM proteins canonically associated with fibrosis (e.g., collagens), several other degraded ECM proteins were present, especially after 28 days of recovery. A limitation of degradome analysis is that the original source of the parent protein cannot be completely verified. However, the CCl_4_ model is relatively specific for liver injury and the majority of the significantly changed peptides were hepatic by Brenda Tissue Ontology analysis (BTO:0000759; *p*=2.2×10^−14^).

One of the most robust families of degraded proteins that were enriched 28 days after cessation of CCl_4_ were associated with the coagulation cascade, especially the fibrin(ogens) (Figure 6, Table 1 and Supplemental Table 4). Previous studies have investigated the potential role of fibrin(ogens) in the development of hepatic injury and fibrosis, with somewhat conflicting results (47, 48). However, few if any studies have investigated the role of fibrin(ogen) ECM in resolution of hepatic fibrosis. In contrast, the turnover of fibrin(ogen) ECM has been heavily studied in other diseases of remodeling and recovery. For example, fibrin(ogen) ECM plays a key role in sub-cutaneous wound-healing (49), and is the rationale for the delivery fibrin-enriched products to promote wound healing (50). However, the clearance of the fibrin-containing provisional matrix is also a key step in late stages of wound-healing (49). Since hepatic fibrosis and subcutaneous wound-healing qualitatively and quantitatively share many similarities, it is possible this increase in fibrin(ogen) fragments during the late stage of recovery from hepatic fibrosis represents a similar clearance of fibrin ECM from the provisional matrix. Future studies should specifically investigate this possibility, especially given the use of anticoagulants in fibrotic liver disease and cirrhosis.

As mentioned in the Introduction, collagenous ECM accumulation has been the major focus in the field of hepatic fibrosis (2). Likewise, proteases involved in resolution of collagenous ECM (e.g., MMP2 and MMP9) have also received a sizeable amount of attention (38). Once the technical limitations to identifying peptide fragments of degraded proteins informatically was overcome (44), the fragment sequence and putative cleavage site may yield new information on proteases involved. The predicted protease that generated each peptide was therefore determined by in silico analysis of the cleavage site using the online open-source tool, Proteasix (www.proteasix.org) (29). Although several proteases with established roles in fibrosis and/or resolution were identified (e.g., MMP2), several novel players were predicted by the algorithm.

The most common predicted protease both 1 and 28 d after cessation of CCl_4_ was calpain1 (Capn1) (Figure 7). It should be noted that Capn1 and its isozyme Capn2 cannot be delineated by substrate or activity assays, and expression of both followed a similar pattern in liver after recovery from fibrosis. This finding was validated by investigating levels of the intrinsic substrate for Capn1/2 (α-fodrin) and via an enzymatic assay (Figure 7B). The calpain family are cysteine proteases that cleave a myriad of protein substrates. Although they are generally localized intracellularly, they can be excreted when proteolytically activated (51). Calpain activation has been identified as a key driver of inflammation in other diseases, such as atherosclerosis and cardiac disease (51, 52). Moreover, calpain activation has been identified to be involved in diseases of remodeling (i.e., fibrosis) via mechanisms that remain unclear (53). Interestingly, the predicted substrates for calpain activation not only included key ECM proteins [e.g., collagens and fibrin(ogens)], but also nuclear proteins, such as histones (Figure 7C), which is in-line with previous findings in other organs (54) and the liver (30). Few studies have identified calpain activation to play a role in hepatic injury. However, a recent study indicated that CAPN2 is induced in human livers in HBV-induced fibrosis (30), and it was demonstrated here that CAPN2 expression correlated with the severity of fibrosis in NASH (Figure 7C). The mechanisms by which CAPN2 may mediate these effects are unclear, but work in other organ systems suggest a role in not only ECM metabolism, but also in organelle damage [e.g., (51)]. These data suggest that the role of calpains in liver disease and fibrosis may be underappreciated at this time.

Taken together, the results of this study indicate that remodeling of the liver during recovery from fibrosis is a complex and highly coordinated process that extends well beyond the degradation of the collagenous scar. A key role of MMP12, and by extension elastin, was demonstrated, which is in-line with previous studies. Moreover, it was shown that even during late stages of recovery (i.e., 28 D after CCl_4_ cessation), there is significant enhanced turnover of proteins. These results also indicate that analysis of the plasma degradome may yield new insight into the mechanisms of fibrosis recovery, and by extension, new “theragnostic” targets. Lastly, a novel potential role for calpain activation in the degradation and turnover of proteins was identified.

## Abbreviations

CCl_4_: carbon tetrachloride
ECM: extracellular matrix
ELISA: enzyme-linked immunosorbent assay
GO: gene ontology
LC-MS/MS: liquid Chromatography with tandem mass spectrometry
MMP: matrix metalloprotease
MPM: multiphoton microscopy
rtPCR: reverse-transcriptase polymerase chain reaction
SDS: sodium dodecyl sulfate

## Acknowledgments

Financial support: Supported, in part, by grants from NIH (R01 AA021978, P20 GM113226, P30 DK120531). Pittsburgh Liver Research Center (P30 DK120531) provided imaging through the Advanced Cell and Tissue Imaging Core, housed in the Center for Biologic Imaging. Equipment grants from NIH (S10 RR025676 and S10 OD019973; PI S. Watkins) were used to purchase microscopes used in this study.

