## Supplemental Materials for "Fibrosis resolution in the mouse liver: role of Mmp12 and potential role of Calpain 1/2"

[1]Department of Medicine, Division of Gastroenterology, Hepatology and Nutrition, [2]Department of Pharmacology and Chemical Biology, [3]Department of Surgery, [4]Department of Pathology, [5]Pittsburgh Liver Research Center,[6] Department of Environmental and Occupational Health, University of Pittsburgh, Pittsburgh, PA 15213. [7]Department of Pharmacology and Toxicology, [8]Department of Medicine, Division of Nephrology and Hypertension and [9]University of Louisville Alcohol Research Center, University of Louisville, Louisville, KY 40292

Send all correspondence to: Gavin E. Arteel, PhD, FAASLD
Thomas E. Starzl Biomedical Science Tower
West 1143
200 Lothrop Street
Pittsburgh, PA 15213


### Supplemental Experimental Procedures

Information on rtPCR primers and probes, drugs and chemical assays and antibodies used in this study are summarized in supplemental Tables 1-3.

### Supplemental Results

Supplemental Table 4 lists all peptides significantly increased in mouse plasma 1D and/or 28D after cessation of $CCl_4$, as described in Experimental Procedures, which is shown graphically in Figure 5.

**Supplemental Table 1-Product information for primers used in RT-PCR**

| Gene name | Supplier | Cat. No. |
|---|---|---|
| *Col1a1* | Thermofisher | Mm00801666_g1 |
| *Acta2* | Thermofisher | Mm01546133_m1 |
| *Eln* | Thermofisher | Mm00514670_m1 |
| *Lox* | Thermofisher | Mm00495386_m1 |
| *Loxl2* | Thermofisher | Mm00804740_m1 |
| *P4htm* | Thermofisher | Mm00512331_m1 |
| *Mmp2* | Thermofisher | Mm00439498_m1 |
| *Mmp9* | Thermofisher | Mm00442991_m1 |
| *Timp1* | Thermofisher | Mm01341361_m1 |
| *Timp2* | Thermofisher | Mm00441825_m1 |
| *Timp3* | Thermofisher | Mm00441826_m1 |
| *Mmp8* | Thermofisher | Mm00439509_m1 |
| *Mmp12* | Thermofisher | Mm00500554_m1 |
| *Mmp13* | Thermofisher | Mm00439491_m1 |
| *Mmp14* | Thermofisher | Mm00485054_m1 |
| *Capn1* | Thermofisher | Mm00482964_m1 |
| *Capn2* | Thermofisher | Mm00486669_m1 |

**Supplemental Table 2- Drugs and Chemical assays used in this study**

| Category | Name | Supplier | Cat No. |
|---|---|---|---|
| *Drug/chemicals* | MMP408 | Sigma | 444291 |
| | Purified MMP-12 | AnaSpec | 55525-1 |
| | SUC-LEU-LEU-VAL-TYR-AMC | Fisher Scientific | 501017021 |
| *Chemical assays* | AST assay kit | Thermofisher | TR70121 |
| | ALT assay kit | Thermofisher | TR71121 |
| | MMP-12 activity assay kit | AnaSpec | AS-71157 |
| | Mouse Desmosine ELISA Kit | MyBioSource Inc | MBS754634 |

**Supplemental Table 3- Primary antibodies used in this study**

| Antibody | Supplier | Cat No. |
|---|---|---|
| GAPDH | CELL SIGNALING | 5174S |
| Elastin | ABCAM | ab21607 |
| α-Fodrin | CELL SIGNALING | 2122S |

### Supplemental Table 4-Plasma peptides increased 1D and/or 28D after cessation of $CCl_4$

| Parent Protein* | | | | | | Predicted Protease (PPV/NPV) | |
|---|---|---|---|---|---|---|---|
| Gene name | Accession number | Peptide sequence | MW (Da) | Log2 FC | -Log10 PV | N-terminus | C-terminus |
| **Increased 1D after $CCl_4$, Decreased 28D after $CCl_4$ (1D data are summarized)** | | | | | | | |
| HBA | P01942 | K.IGGHGAEYGAEALERM.F | 1936.19 | 1.55 | 2.30 | N/D | ADAMTS4(94/85) |
| H12 | P15864 | L.KKALAAAGYDVEKNNSRIKLG.L | 2472.94 | 1.49 | 1.80 | ADAMTS4(94/85) | Capn1(84/58) |
| | | K.ALAAAGYDVEKNNSRIKLGL.K | 2359.77 | 1.19 | 2.27 | | |
| APOC1 | P34928 | P.DLSGTLESIPDK.L | 1484.68 | 2.86 | 2.28 | ADAMTS4(94/85) | ADAMTS4(94/85) |
| H15 | P43276 | A.PAETAAPAPVEKSPAKKK(+42.02)T(-.98).T | 2093.42 | 1.06 | 2.02 | Capn1(84/58) | Capn1(84/58) |
| H13 | P43277 | A.AAGKRKASGPPVSELIT.K | 1881.22 | 1.10 | 1.59 | Capn1(84/58) | Capn1(84/58) |
| HMGB1 | P63158 | K.YEKDIAAYRAKGKPDAA.K | 2123.46 | 1.57 | 2.32 | Capn1(84/58) | ADAMTS4(94/85) |
| | | R.YEREM(+15.99)KTYIPPKGETKKKFKDPNAPKRPPSAF.F | 4083.81 | 1.06 | 1.52 | Try3(78/93) | N/D |
| H2AV | Q3THW5 | I.RGDEELDSLI.K | 1387.56 | 1.22 | 2.60 | ELANE(77/78) | Capn1(84/58) |
| CO5A2 | Q3U962 | G.QRGAPGKDGEVGPSG.P | 1565.67 | 2.21 | 2.25 | ADAMTS4(94/85) | Capn1(84/58) |
| | | P.F(+43.01)GP(+15.99)R(+.98)GPPGPVGPSGKEGNPG.P | 2055.30 | 1.20 | 1.58 | N/D | Ctsb(79/81) |
| TAGL2 | Q9WVA4 | G.DPNWFPK.K | 1088.24 | 1.19 | 1.53 | Mep1a(78/66) | N/D |
| **Increased 1D after $CCl_4$, not significantly changed 28D after $CCl_4$ (1D data are summarized)** | | | | | | | |
| IMDH2 | P24547 | D.AGVDALRV.G | 972.07 | 6.65 | 1.39 | Casp1(95/96) | Ctsb(79/81) |
| H11 | P43275 | A.GYDVEKNNSRIKLGL.K | 1905.20 | 8.75 | 1.93 | Ctsb(79/81) | Capn1(84/58) |
| CEBPA | P53566-3 | S.RQQEKAKAAAGPAG.G | 1526.68 | 11.50 | 1.40 | N/D | ADAMTS4(94/85) |
| H2A2A | Q6GSS7 | L.PKKTESHHKAKG.K | 1588.88 | 4.47 | 1.68 | Capn1(84/58) | N/D |
| **Decreased 1D after $CCl_4$, increased 28D after $CCl_4$ (28D data are summarized)** | | | | | | | |
| FIBA | E9PV24-2 | T.D(+42.01)(+15.99)TED(-18.01)KGEFLSEGGGVR.G | 1853.93 | 4.38 | 1.33 | N/D | capn1(84/58) |
| | | T.NIEDPSSHVPEF.S | 1558.64 | 1.69 | 3.12 | ELANE(77/78) | PGC(100/100) |
| ARI1B | E9Q4N7 | Y.SRPGAGGGGGGGGGGGSGGGGGGGAGGAGGAAAA AAGAGAVAAAA.A | 3385.46 | 2.61 | 1.90 | capn1(84/58) | Mcpt3(93/95) |
| ECH1 | O35459 | T.SGIDLM(+15.99)DM(+15.99).A | 1053.23 | 5.21 | 3.41 | ELANE(77/78) | CTSD(79/85) |
| WDR1 | O88342 | T.GHNKVIN.S | 969.07 | 4.24 | 1.45 | ADAMTS4(94/85) | Mcpt3(93/95) |
| CO3 | P01027 | L.W(+15.99)ENGNLL.R | 1114.28 | 4.31 | 5.31 | CTSD(79/85) | ADAMTS4(94/85) |

| Parent Protein* | | | | | | Predicted Protease (PPV/NPV) | |
|---|---|---|---|---|---|---|---|
| Gene name | Accession number | Peptide sequence | MW (Da) | Log2 FC | -Log10 PV | N-terminus | C-terminus |
| SPA3K | P07759 | T.SAQSILFM(+15.99)AKVNNPK | 1749.08 | 2.79 | 2.11 | Ctsb(79/81) | N/D |
| APOE | P08226 | N.PIITPVAQENQ | 1323.48 | 3.19 | 5.69 | capn1(84/58) | N/D |
| APOA2 | P09813 | R.Q(-17.03)ADGPDM(+15.99)QSLFTQY.F | 1904.11 | 2.61 | 1.31 | capn1(84/58) | ADAMTS4(94/85) |
| H12 | P15864 | R.RKASGPPVSELITKAVAASKE.R | 2451.88 | 2.88 | 1.65 | TMPRSS11D(--/99) | ADAMTS4(94/85) |
| | | K.ASGPPVSEL.I | 1097.29 | 4.83 | 2.30 | capn1(84/58) | CTSD(79/85) |
| GSTP1 | P19157 | N.QGGKAFIV.G | 990.14 | 2.85 | 1.74 | ADAMTS4(94/85) | ELANE(77/78) |
| CATA | P24270 | G.YGSHTFK.L | 1009.14 | 5.73 | 2.31 | N/D | capn1(84/58) |
| H15 | P43276 | S.KGTLVQTKGTGASGSFKLN.K | 2109.43 | 4.25 | 3.30 | capn1(84/58) | capn1(84/58) |
| | | G.TLVQTKGTGASGSFKLN.K | 1894.17 | 3.69 | 2.62 | | |
| MAF | P54843 | H.PTAGAPGAAGGASAS.A | 1350.42 | 3.21 | 1.43 | N/D | capn1(84/58) |
| H2B1A | P70696 | P.GELAKHAVSEGTKAVTKYTSS.K | 2389.71 | 1.30 | 2.43 | Casp1(95/96) | N/D |
| CO5A2 | Q3U962 | P.RGLVGPPGS.R | 1092.28 | 1.81 | 2.54 | ADAMTS4(94/85) | capn1(84/58) |
| MARH6 | Q6ZQ89-3 | N.NNAVPAGEGL.H | 1192.26 | 3.18 | 2.32 | ADAMTS4(94/85) | ELANE(77/78) |
| NPNT | Q91V88 | Y.LTVSAAKAPGGKAAR.L | 1673.99 | 3.01 | 1.32 | Mcpt3(93/95) | capn1(84/58) |
| BHMT2 | Q91WS4 | E.VGAPVAVTM(+15.99).C | 1076.31 | 5.07 | 3.23 | capn1(84/58) | Ctsb(79/81) |
| HEMO | Q91X72 | V.ASPLPT(-18.01)ANGRVAEVE.N | 1723.92 | 5.13 | 3.07 | capn1(84/58) | capn1(84/58) |
| PARD3 | Q99NH2-5 | K.KGTEGLGF.S | 1023.16 | 3.45 | 3.50 | capn1(84/58) | Cma1(93/95) |
| **Not significantly changed 1D after $CCl_4$, increased 28D after $CCl_4$ (2D data are summarized)** | | | | | | | |
| FIBA | E9PV24-2 | T.TDT(-18.01)EDKGEFLS(-18.01)EGGGVR.G | 1955.04 | 2.59 | 1.64 | N/D | capn1(84/58) |
| | | T.TDTED(+15.99)KGEFLSEGGGVR.G | 1955.04 | 2.09 | 1.34 | | |
| | | T.TDTEDKGEFLS(-18.01)EGGGVR.G | 1955.04 | 2.26 | 1.61 | | |
| | | T.TDTED(-18.01)KGEFLS(-18.01)EG.G | 1585.61 | 2.93 | 1.59 | | Mcpt3(93/95) |
| | | T.TDTEDKGEFLS(-18.01)EG.G | 1585.61 | 2.62 | 1.54 | | |
| | | T.TDTEDKGEFLS(-18.01)E.G | 1528.56 | 2.26 | 1.53 | | Casp1(95/96) |
| | | T.TDT(-18.01)EDKGEFLS(-18.01).E | 1471.50 | 2.51 | 1.55 | | ADAMTS4(94/85) |
| | | T.TDTEDKGEFLS(-18.01).E | 1471.50 | 2.26 | 1.57 | | |
| | | T.D(-18.01)TEDKGEFLSEGGGVR.G | 1853.93 | 2.33 | 1.62 | | capn1(84/58) |

| Parent Protein* | | | | | | Predicted Protease (PPV/NPV) | |
|---|---|---|---|---|---|---|---|
| Gene name | Accession number | Peptide sequence | MW (Da) | Log2 FC | -Log10 PV | N-terminus | C-terminus |
| | | D.TEDKGEFLS(-18.01)EGGGVR.G | 1752.82 | 2.31 | 1.41 | | |
| | | D.TEDKGEFLSEGGGVR.G | 1752.82 | 1.85 | 1.44 | | |
| ARI1B | E9Q4N7 | G.G(+27.99)GGGGGGGGGS(-18.01)GGGGGGGG.A | 1260.16 | 3.17 | 4.85 | ADAMTS4(94/85) | ADAMTS4(94/85) |
| PSB1 | O09061 | P.VGSYQRDSF.K | 1283.42 | 10.65 | 1.45 | Mcpt3(93/95) | N/D |
| ILK | O55222 | H.SGIDFKQL.N | 1158.29 | 8.42 | 2.04 | Mep1a(78/66) | CTSD(79/85) |
| ADH1 | P00329 | W.KGAIFGGF.K | 1110.33 | 2.60 | 1.44 | capn1(84/58) | capn1(84/58) |
| CO3 | P01027 | F.RLLW(+15.99)ENGNLL.R | 1530.81 | 1.35 | 1.66 | Cma1(93/95) | ADAMTS4(94/85) |
| | | R.LLW(+15.99)ENGNLL.R | 1383.63 | 6.23 | 1.82 | KLK6(80/96) | |
| | | R.LLWENGNLL.R | 1383.63 | 1.29 | 1.39 | | |
| | | L.LWENGNLL.R | 1227.44 | 3.84 | 2.70 | N/D | |
| | | L.WENGNLL.R | 1114.28 | 3.04 | 3.01 | CTSD(79/85) | |
| H12 | P15864 | A.AAVTKKVAKSPK.K | 1426.78 | 8.99 | 3.19 | capn1(84/58) | capn1(84/58) |
| | | R.KASGPPVSELITKAVAA.S | 1882.21 | 9.04 | 1.33 | TMPRSS11D(--/99) | capn1(84/58) |
| PH4H | P16331 | Q.NGAVSLIF.S | 1035.18 | 2.53 | 1.32 | N/D | capn1(84/58) |
| VIME | P20152 | L.LIKTVETRDGQVINETSQHHDDLE | 2891.17 | 12.92 | 1.40 | capn1(84/58) | N/D |
| CATA | P24270 | G.YGSHTFKL.V | 1108.27 | 3.98 | 2.77 | N/D | CTSD(79/85) |
| | | G.YGSHT(-18.01)FK.L | 1009.14 | 12.22 | 3.70 | | capn1(84/58) |
| | | Y.GSHT(-18.01)FKL.V | 1051.22 | 6.56 | 2.18 | | |
| URIC | P25688 | L.RGIRNIETF.A | 1289.51 | 3.30 | 2.39 | capn1(84/58) | Mcpt3(93/95) |
| ACOC | P28271 | F.TGVPAVVDF.A | 1122.30 | 3.28 | 1.37 | capn1(84/58) | capn1(84/58) |
| NOTC4 | P31695 | P.CLNGGSCSIRPEGYSCTC.L | 2060.40 | 2.29 | 3.10 | N/D | N/D |
| NLTP | P32020 | V.VGVGM(+15.99)TKF.M | 1068.38 | 3.03 | 3.01 | capn1(84/58) | capn1(84/58) |
| | | I.GGIFAFKV.K | 1079.36 | 3.44 | 3.59 | ELANE(77/78) | ELANE(77/78) |
| H14 | P43274 | A.PAAPAAPAPAEKTPVKKKARKAAGGAKR.K | 2938.54 | 4.76 | 7.75 | capn1(84/58) | TMPRSS11D(--/99) |
| ELN | P54320 | G.ALGGLVPGAVPGALPGAVPAVPGAGGVPGAGTPAAAAAAAAK.A | 3598.19 | 3.06 | 5.12 | TMPRSS11D(--/99) | capn1(84/58) |
| YBOX1 | P62960 | R.E(-18.01)DGNEEDKENQGDETQGQQPPQRRY.R | 3260.34 | 12.00 | 2.24 | KLK6(80/96) | ADAMTS4(94/85) |
| UFD1 | P70362 | K.RGIPNYEF.K | 1251.46 | 3.22 | 2.40 | capn1(84/58) | CTSD(79/85) |

| Parent Protein* | | | | | | Predicted Protease (PPV/NPV) | |
|---|---|---|---|---|---|---|---|
| Gene name | Accession number | Peptide sequence | MW (Da) | Log2 FC | -Log10 PV | N-terminus | C-terminus |
| FA7 | P70375 | D.RGATALELM(+15.99).S | 1163.32 | 11.67 | 2.59 | ADAMTS4(94/85) | ADAMTS4(94/85) |
| H2B1A | P70696 | K.HAVSEGTKAVTKYT.S | 1706.93 | 2.07 | 2.34 | capn1(84/58) | N/D |
| CO1A2 | Q01149 | G.SPGEAGSAGPAGP.P | 1208.26 | 3.00 | 1.52 | capn1(84/58) | N/D |
| PYC | Q05920 | G.NGALFVEKF.I | 1194.41 | 4.09 | 1.38 | TMPRSS11D(--/99) | capn1(84/58) |
| MAST2 | Q60592 | K.LAAALAAAEKK.L | 1297.61 | 1.55 | 1.88 | ADAMTS4(94/85) | capn1(84/58) |
| NCK5L | Q6GQX2 | K.KAQILEV.L | 1041.31 | 3.07 | 1.77 | capn1(84/58) | N/D |
| SAM50 | Q8BGH2 | D.RGVSAEYSF.P | 1227.31 | 3.38 | 3.19 | capn1(84/58) | N/D |
| WIPF1 | Q8K1I7 | G.G(+27.99)GGGGGGGGGGS(-18.01)GGN.F | 1222.16 | 2.66 | 3.03 | ADAMTS4(94/85) | ADAMTS4(94/85) |
| ATTY | Q8QZR1 | M.VGIEM(+15.99)EHF.P | 1189.43 | 2.63 | 3.31 | N/D | N/D |
| AAMDC | Q8R0P4 | K.QGIDVRVL.Q | 1155.37 | 7.07 | 5.39 | Try3(78/93) | CTSD(79/85) |
| | | A.QGVRVGGVF.H | 1126.29 | 9.91 | 1.56 | Cma1(93/95) | N/D |
| KPRB | Q8R574 | K.AGLTHLITM(+15.99).D | 1199.45 | 4.45 | 2.28 | Gzma(80/95) | N/D |
| ES8L3 | Q91WL0 | W.LVKNEAGLTG.Y | 1350.55 | 2.07 | 2.98 | N/D | capn1(84/58) |
| BHMT2 | Q91WS4 | R.AGADVLQTF.T | 1178.32 | 3.19 | 1.86 | ADAMTS4(94/85) | Mcpt3(93/95) |
| METK1 | Q91X83 | N.EDITLEAM(+15.99).Q | 1163.28 | 4.86 | 4.77 | N/D | ADAMTS4(94/85) |
| THIKA | Q921H8 | C.TGARQVVTLL.N | 1274.52 | 3.45 | 2.91 | capn1(84/58) | capn1(84/58) |
| DCTN2 | Q99KJ8 | R.VGTKGLDF.S | 1079.23 | 3.99 | 1.87 | capn1(84/58) | capn1(84/58) |
| TM218 | Q9CQ44 | V.LGVGAGVFLL.A | 1115.39 | 1.72 | 2.62 | capn1(84/58) | Mep1a(78/66) |
| NNRD | Q9CZ42-2 | K.VGADLTHVF.C | 1189.41 | 1.66 | 1.55 | ADAMTS4(94/85) | N/D |
| ZN687 | Q9D2D7 | A.VPPVPGPLALPV.L | 1339.70 | 1.18 | 1.71 | capn1(84/58) | ADAMTS4(94/85) |
| D42E1 | Q9D665 | K.TGVTHYFSL.E | 1281.44 | 8.26 | 1.36 | capn1(84/58) | ADAMTS4(94/85) |
| NBEA | Q9EPN1-3 | A.GGGAGGGGAMGEPRGAAGSGPVV.L | 2010.23 | 16.23 | 1.47 | capn1(84/58) | ADAMTS4(94/85) |
| SCN4A | Q9ER60 | L.IKIIGNSVGALG.N | 1368.65 | 1.26 | 1.54 | CTSD(79/85) | Mcpt3(93/95) |
| PYGL | Q9ET01 | I.VGVENVAEL.K | 1170.38 | 4.54 | 3.53 | ELANE(77/78) | ADAMTS4(94/85) |
| **Increased 1D after $CCl_4$, increased 28D after $CCl_4$ (28D data are summarized)** | | | | | | | |
| TITIN | A2ASS6 | A.INAAGVGPASLPSDPVTARDP.V | 2175.45 | 5.08 | 1.65 | Ctsb(79/81) | capn1(84/58) |

| Parent Protein* | | | | | | Predicted Protease (PPV/NPV) | |
|---|---|---|---|---|---|---|---|
| Gene name | Accession number | Peptide sequence | MW (Da) | Log2 FC | -Log10 PV | N-terminus | C-terminus |
| FIBA | E9PV24-2 | T.TDTE(+21.98)DKGEFLSEGGGVR.G | 1955.04 | 4.06 | 1.33 | N/D | capn1(84/58) |
| | | T.DTEDKGEFLSEGGGVR.G | 1853.93 | 2.56 | 1.47 | | |
| | | T.DTEDKGEFLS(-18.01)EG.G | 1484.50 | 12.80 | 1.42 | | Mcpt3(93/95) |
| KI67 | E9PVX6 | S.RGRDAGTPAPMQEGNGTTAIMET.P | 2545.81 | 4.95 | 1.31 | capn1(84/58) | N/D |
| ECH1 | O35459 | L.RGTSQLYF.N | 1198.36 | 5.77 | 7.65 | capn1(84/58) | Cma1(93/95) |
| | | D.SGLVSRVF.Q | 1107.24 | 4.78 | 3.40 | Casp1(95/96) | capn1(84/58) |
| CO3 | P01027 | F.R(+15.99)LLW(+15.99)ENGNLL.R | 1530.81 | 3.15 | 2.00 | Mcpt3(93/95) | ADAMTS4(94/85) |
| | | F.RLLWENGNL.L | 1374.62 | 2.11 | 1.67 | | |
| | | F.RLLWENG.N | 1148.30 | 10.97 | 5.66 | | N/D |
| | | L.LW(+15.99)ENGNLL.R | 1227.44 | 5.69 | 3.49 | N/D | ADAMTS4(94/85) |
| SPA3K | P07759 | A.QSILFM(+15.99)AKVNNPK | 1560.89 | 10.54 | 1.44 | capn1(84/58) | N/D |
| APOA2 | P09813 | R.Q(-17.03)ADGPDM(+15.99)QSL.F | 1364.51 | 8.78 | 1.87 | capn1(84/58) | CTSD(79/85) |
| CO1A1 | P11087 | R.P(+27.99)GEAGLPGAK(+27.99)GLTGS(+21.98).P | 1564.77 | 13.27 | 1.60 | capn1(84/58) | capn1(84/58) |
| | | G.PP(+15.99)GATGFP(+15.99)GAAGRVGPP(+15.99)GP.S | 1804.01 | 3.54 | 1.91 | capn1(84/58) | ADAMTS4(94/85) |
| PH4H | P16331 | D.IGATVHEL.S | 1041.13 | 4.07 | 5.02 | CTSD(79/85) | N/D |
| ENOA | P17182 | R.IGAEVYHNL.K | 1299.51 | 10.82 | 2.13 | capn1(84/58) | N/D |
| CATA | P24270 | A.KGAGAFGYF.E | 1117.23 | 4.97 | 3.92 | capn1(84/58) | PGC(100/100) |
| MUG1 | P28665 | S.TGSFSQKF.Q | 1116.21 | 14.44 | 1.47 | capn1(84/58) | N/D |
| NLTP | P32020 | S.TGSTALFM(+15.99).A | 985.13 | 3.74 | 5.35 | N/D | Ctsb(79/81) |
| APOC3 | P33622 | F.WDSNPEDQPTPA.I | 1616.72 | 7.38 | 1.79 | N/D | N/D |
| HPPD | P49429 | G.AGVQHIAL.K | 993.18 | 4.30 | 2.80 | Mep1a(78/66) | CTSD(79/85) |
| HXK4 | P52792 | T.VGVDGSVYKL.H | 1274.45 | 14.70 | 4.28 | Ctsb(79/81) | N/D |
| OSTR | P54615 | E.SDKAFM(+15.99)SKQEGNKVVNRL.R | 2336.68 | 14.28 | 4.28 | capn1(84/58) | ADAMTS4(94/85) |
| TBA1A | P68369 | R.LDHKFDLM(+15.99).Y | 1337.58 | 11.33 | 3.16 | MMP14(75/81) | N/D |
| GSK3A | Q2NL51 | G.GGGGPGGSASGPGGTGGGKASVGAMGGGVGASSS.G | 2661.78 | 14.39 | 1.38 | ADAMTS4(94/85) | Casp1(95/96) |
| AP4AT | Q3U3N6 | H.THPNPGNLL.P | 1196.34 | 7.87 | 6.51 | capn1(84/58) | capn1(84/58) |
| NEUL4 | Q5NCX5-2 | P.GGGGGPGSSGPGLGSGGGLGGGGELHP.R | 2342.49 | 3.39 | 1.49 | ADAMTS4(94/85) | N/D |
| POSTN | Q62009-5 | A.ITGGAVFET.M | 1096.28 | 10.17 | 1.35 | CTSD(79/85) | ADAMTS4(94/85) |

| Parent Protein* | | | | | | Predicted Protease (PPV/NPV) | |
|---|---|---|---|---|---|---|---|
| Gene name | Accession number | Peptide sequence | MW (Da) | Log2 FC | -Log10 PV | N-terminus | C-terminus |
| DDX3Y | Q62095 | K.KGADSLENF.L | 1221.39 | 11.30 | 1.48 | capn1(84/58) | capn1(84/58) |
| ZEB1 | Q64318 | S.ITDAADCEGGMPD.D | 1496.56 | 9.86 | 1.38 | CTSD(79/85) | GZMB(78/97) |
| CP2CT | Q64458 | P.KGTTVITSL.S | 1103.29 | 2.79 | 1.68 | N/D | Mcpt3(93/95) |
| MARH6 | Q6ZQ89-3 | A.NGIRNIDL.H | 1122.26 | 4.85 | 2.01 | N/D | N/D |
| SI1L2 | Q80TE4 | S.GLVSIKAF.Y | 1084.29 | 4.35 | 2.29 | capn1(84/58) | Mcpt3(93/95) |
| PAQR2 | Q8BQS5 | F.HGVSNLQEF.R | 1333.48 | 3.77 | 3.13 | N/D | N/D |
| NAKD2 | Q8C5H8-4 | G.TGSKAWSF.N | 1054.14 | 3.17 | 2.49 | N/D | capn1(84/58) |
| THIKB | Q8VCH0 | C.TGARQVVTL.L | 1160.41 | 3.71 | 2.78 | capn1(84/58) | PGC(100/100) |
| THIKA | Q921H8 | C.TGARQVVTL.L | 1160.41 | 3.71 | 2.78 | N/D | N/D |
| SNTB1 | Q99L88 | K.QGIETHLF.R | 1228.43 | 2.63 | 1.69 | capn1(84/58) | capn1(84/58) |
| BI1 | Q9D2C7 | A.AGAYVHVV.T | 987.13 | 3.92 | 1.97 | capn1(84/58) | ELANE(77/78) |
| STR6L | Q9DBN1-2 | H.RGLEM(+15.99)LQGF.G | 1244.45 | 9.67 | 1.70 | N/D | N/D |
| TIM23 | Q9WTQ8 | M.KGSLLQQSL | 1104.35 | 11.37 | 1.99 | N/D | N/D |
| PDCD7 | Q9WTY1 | A.EASRGGGGGGAFFP.V | 1436.56 | 10.37 | 1.58 | Mcpt3(93/95) | capn1(84/58) |
| DECR2 | Q9WV68 | P.NGIKQLLEF.E | 1287.49 | 1.62 | 1.60 | N/D | Mcpt3(93/95) |

Log2FC and -Log10PV are in comparison to values from control mice. Periods between amino acids indicates the detected cleavage site. Red parent proteins are defined as "Matrisome" by the matrisome project (www.matrisomeproject.mit.edu) (2). The predicted protease output for N-terminus and C-terminus of the peptide fragment is derived from output from ProteaSix analysis (3); as are the positive- (PPV) and negaaretive- (NPV) predictive values. See also Figures 5 and 6. Mass addition (e.g., +42.02) and mass loss (e.g., -18.01) denotes chemical modification of the amino acid, corresponding to the following:

| | |
|---|---|
| -18.01 | dehydration |
| -17.03 | pyroglutamic acid |
| -0.98 | amidation |
| +0.98 | deamidation |
| +15.99 | oxidation |
| +18.01 | hydration |
| +21.98 | sodiation |
| +27.99 | formylation |
| +28.03 | dimethylation |

| | |
|---|---|
| +42.01 | acetylation |
| +42.02 | guanidiation |
| +43.01 | carbamylation |
| +79.97 | phosphorylation |
| +156.12 | 4-hydroxynonenal |
| +162.05 | hexose |

*Some peptides mapped to homologous locations on several protein isoforms. Canonical and/or most abundant parent protein is listed.

N/D: predicted protease not detectable.
